# Sirt5 preserves cardiac function in ischemia-reperfusion injury by inhibiting ANT2 lactylation

**DOI:** 10.1101/2024.11.25.625148

**Authors:** Simeng Li, Siman Shen, Yongfang Hong, Kun Ding, Suyun Chen, Jianning Chen, Changsen Wang, Yaofeng Wen, Guixi Mo, Lili Yu, De-Li Shi, Liangqing Zhang

**Affiliations:** Department of Anesthesiology, the Second Affiliated Hospital of Guangdong Medical University, Zhanjiang, China; Faculty of Chinese Medicine, The State Key Laboratory for Quality Research in Chinese Medicines of the Macau University of Science and Technology; Department of Cardiovascular Internal Medicine, the Second Affiliated Hospital of Guangdong Medical University, Zhanjiang, China; Department of Anesthesiology, Affiliated Hospital of Guangdong Medical University, Zhanjiang, China; Laboratory of Developmental Biology, CNRS-UMR7622, Institut de Biologie Paris-Seine (IBPS), Sorbonne University, Paris, France

**Author notes:** Equal contributions (Simeng Li, Siman Shen, Yongfang Hong). Authors for correspondence: Lili Yu De-Li Shi Liangqing Zhang.

**Keywords:** Sirt5, ANT2, VDAC1, ischemia-reperfusion injury, mitochondrial dysfunction, cardiac protection

## Abstract

Myocardial ischemia-reperfusion (IR) injury is the most common type of heart disease. IR-disrupted mitochondrial homeostasis affects heart energy metabolism and function, but the mechanism remains poorly understood. Here we report a cardioprotective role of the mitochondrial protein Sirt5 in a mouse model of cardiac IR injury. The down-regulation of Sirt5 in IR-injured heart is associated with an increased protein lactylation and impaired mitochondrial function. Overexpression of *Sirt5* alleviates, whereas conditional knockout of *Sirt5* aggravates, mitochondrial damage and cardiac injury. Mechanistically, Sirt5 interacts with the inner mitochondrial membrane protein adenine nucleotide translocase 2 (ANT2), inhibiting its lysine lactylation to promote its association with the outer mitochondrial membrane protein voltage-dependent anion-selective channel 1 (VDAC1). Lactylation-resistant ANT2 efficiently complexes with VDAC1 and improves cardiac function after injury. Therefore, we uncover a Sirt5-regulated interaction of ANT2 and VDAC1 in maintaining mitochondrial homeostasis, highlighting the potential of targeting ANT2 lactylation as a therapeutic strategy in the treatment of cardiac injury.

**Teaser:** Sirt5 promotes the interaction of mitochondrial proteins to protect mitochondrial homeostasis and improve heart function in ischemia-reperfusion injury.

## Introduction

Myocardial ischemia is a common pathogenesis of cardiac dysfunction caused by an absolute reduction of coronary blood flow (*1–3*). Timely restoration of coronary perfusion is crucial for preserving viable myocardial cells around the ischemic lesion, yet it also triggers a second wave of cardiac damage, known as ischemia/reperfusion (I/R) injury (*4*). This can cause an excessive production of reactive oxygen species (ROS) and promote oxidative insult to the heart, leading to myocardial cell dysfunction and death, subsequently affecting the overall function of the heart. As an organ with high energy demands, any abnormality of mitochondrial function can severely impact the oxidative metabolism, which accounts for more than 95% of the energy production in the heart (*5*, *6*). In myocardial IR injury, there is a disruption of energy metabolism, an increased oxidative stress, changes in mitochondrial dynamics, and the activation of mitochondrial autophagy (*7*). Therefore, targeting mitochondrial function represents an emerging treatment strategy in heart failure (*8*).

Sirtuin proteins belong to a highly conserved family of NAD^+^-dependent histone deacetylases that regulate a variety of important signaling pathways. There are seven sirtuin paralogs in mammals, namely Sirt1 to Sirt7. Sirt1 is mainly present in the nucleus but can shuttle to the cytoplasm under specific conditions (*9*, *10*). Sirt2 is primarily found in the cytoplasm but also accumulates in the nucleus during the G2 to M phase transition of the cell cycle (*11*). Sirt3-5 have mitochondrial targeting sequences and are essentially localized to the mitochondria (*12–14*). Sirt6 and Sirt7 are also nuclear proteins, with Sirt6 mostly associated with chromatin (*15*) and Sirt7 enriched in the nucleolus (*16*). Sirtuin proteins have attracted widespread attention due to their key role in regulating histone deacetylation (*17*). Several sirtuin proteins also exert non-histone deacetylase and mono-ADP-ribosyltransferase activities to regulate cell proliferation, cell senescence, DNA damage, oxidative stress, inflammation, and cellular metabolism (*18*, *19*). Evidence is accumulating that sirtuin proteins are critically involved in several post-translational modifications (PTMs) important for cardiac function and represent promising therapeutic targets for the treatment of cardiovascular diseases (*18*, *20*, *21*). Sirt5 displays unique enzymatic activity that can hydrolize negatively charged lysine modifications (*22–24*), while functioning as a very weak deacetylase (*25*). For example, it prevents lysine succinylation to maintain cardiac metabolism in cardiomyopathy and improve cardiac function (*26*, *27*). The loss of Sirt5 in mice with diabetic cardiomyopathy aggravates myocardial injury by increasing lysine malonylation modification and decreasing the stability of glutathione S-transferase P (GSTP1) protein (*28*). However, it is still unclear how Sirt5 regulates other PTMs associated with mitochondrial homeostasis and cardioprotection.

Lactate, an important carbon-containing metabolic product of the cellular glycolytic pathway, emerges as a key energy substrate and signaling molecule. Lactylation is a novel type of protein PTM involved in the metabolic regulation of gene expression and function (*29*). The effects of lactate signaling and protein lactylation in cardiovascular disease remain poorly understood. Elevated intracellular lactate concentrations can promote cardiac regeneration in zebrafish by inducing metabolic switch toward glycolysis (*30*) and alleviate heart failure in mice by modulating sarcomere integrity through lactylation modification of α-myosin heavy chain (*31*). Histone lactylation can trigger gene expression for the cardiac repair process post-myocardial infarction (*32*). However, it has been also reported that an increased lactate level after myocardial infarction leads to endothelial-to-mesynchymal transition by inducing the lactylation of Snail1, thereby promoting cardiac fibrosis and exacerbating heart dysfunction (*33*). In addition, there is evidence that lactic acid promotes pulmonary fibrosis by inducing myofibroblast differentiation through pH-dependent activation of TGFß signaling (*34*), and that a high level of blood lactate is correlated with high mortality in patients with heart failure (*35*). Therefore, lactate-induced protein lactylation modifications may exert context-dependent effects on cardiac function, but how they regulate mitochondrial function following cardiac injury merits further investigation.

In this study, we uncover an essential role of Sirt5 in maintaining cardiac mitochondrial homeostasis by regulating lysine lactylation modification of adenine nucleotide translocase 2 (ANT2), an inner mitochondrial protein. Specifically, Sirt5 interacts with ANT2 to prevent its lactylation modification on lysine 163 caused by IR-induced increase of lactate levels. This promotes the association between ANT2 and voltage dependent anion-selective channel 1 (VDAC1) located on the outer mitochondrial membrane, thereby facilitating the communication of metabolites between the cytosol and mitochondrial matrix and maintaining mitochondrial homeostasis. Overexpression of Sirt5 is able to rescue IR-induced mitochondrial damage and improve cardiac function. Furthermore, the substitution of the lysine residue at position 163 in ANT2 by an arginine residue enhances its interaction with VDAC1 and consequently, reverses mitochondrial dysfunction caused by IR injury or deficiency of Sirt5. These findings provide insights into Sirt5-dependent delactylation in the regulation of energy metabolism and cardiac function. They should help design therapeutic strategy for the intervention of heart failure.

## Results

### Sirt5 protects mitochondrial and cardiac functions in IR injury

Although several Sirt proteins are implicated in heart failure, IR injury and cardiac hypertrophy, the functions of mitochondrial Sirt4 and Sirt5 in these pathological processes remain largely elusive (*21*). Therefore, we first performed western blot analyses to examine changes of Sirt protein levels in mouse ventricular myocardial tissues subjected to 45 minutes of ischemia and 6 hours of reperfusion (IR injury). Sirt1-3 protein levels were up-regulated, whereas Sirt4-7 were down-regulated after IR injury (Fig. 1, A and B). Quantification of Sirt1-7 protein levels indicated that Sirt5 was most strongly decreased in IR-injured myocardium (Fig. 1B). Therefore, we thus focused the study on this protein. Immunohistochemical analysis further confirmed a decreased expression of Sirt5 protein in the left ventricular area of IR-injured mouse heart (Fig. 1C). Importantly, cardiac tissues from patients subjected to IR due to heart valve replacement also displayed reduced SIRT5 protein levels compared to normal heart tissues (Fig. 1D). These observations indicate that cardiac injury negatively regulates the expression of Sirt5 protein, which may be of clinical significance. Moreover, in HL-1 mouse cardiomyocytes subjected to hypoxia for 12 hours and reoxygenation for 4 hours (HR injury), there was also a decreased expression of Sirt5 protein (Fig. 1E). We then confirmed the enrichment of Sirt5 in the mitochondria of HL-1 cells (Fig. 1F) and verified its down-regulation in this organelle following HR injury (Fig. 1G).

**Fig. 1.**
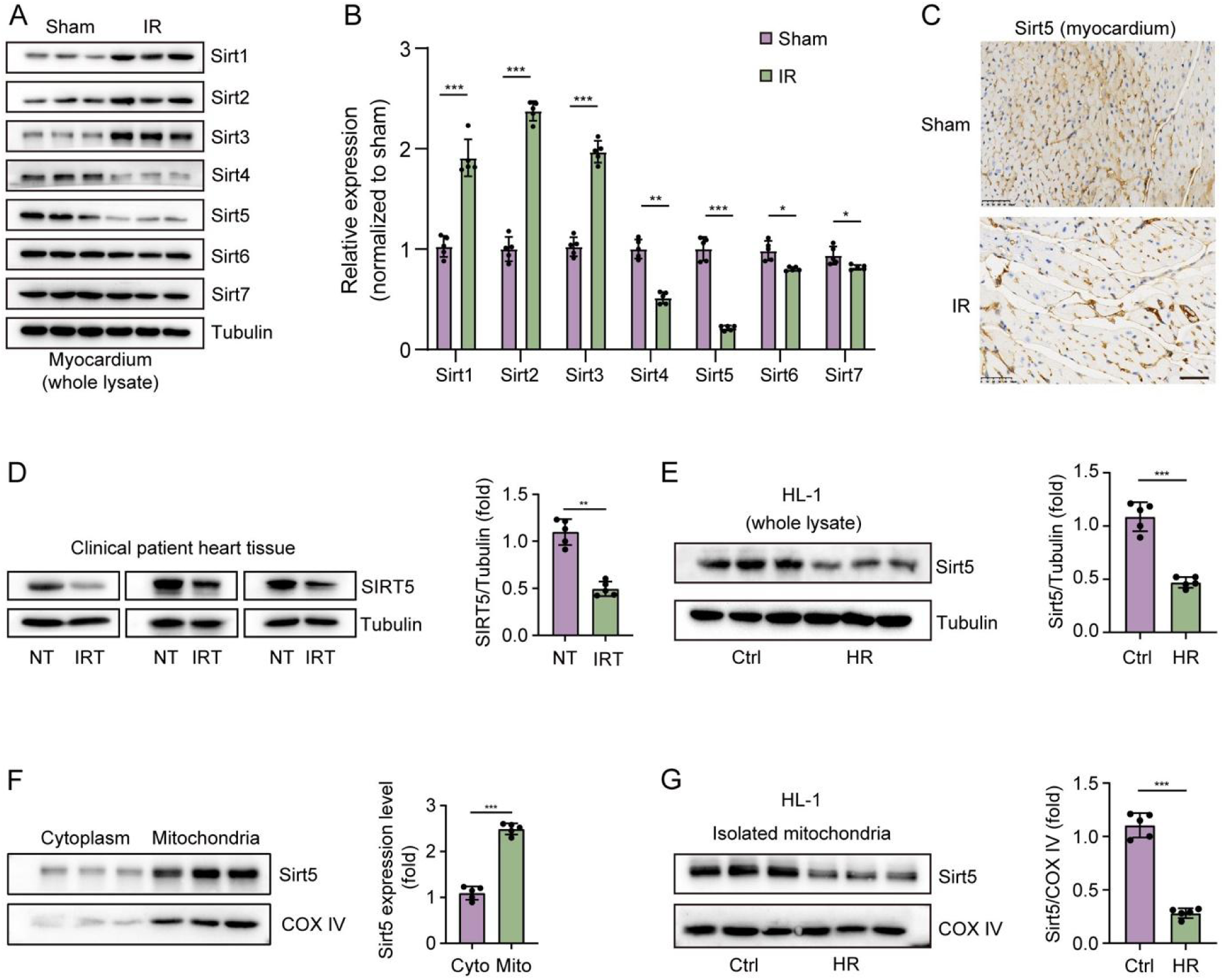
Sirt5 is a mitochondria-enriched protein down-regulated under IR and HR conditions. (**A**) Western blot analysis of Sirt1-7 protein expression in mouse myocardial tissues 6 hours after IR injury. (**B**) Quantification of Sirt1-7 expression in IR-injured myocardium. Protein levels in sham condition are set to 1 after normalization to GAPDH. Two-tailed *t*-test (*, *P* < 0.05; **, *P* < 0.01; ***, *P* < 0.001). (**C**) Representative images of Sirt5 immunohistochemistry in sham and IR conditions. Scale bar, 200 µm. (**D**) Western blot analysis and quantification of SIRT5 protein expression levels in normal tissues (NT) and IR-injured myocardial tissues (IRT) from human patients. (**E**) Western blot analysis and quantification of Sirt5 protein expression levels in HL-1 cells 48 hours after HR injury. (**F**) Western blot analysis and quantification of Sirt5 protein accumulation in the mitochondria. Two-tailed *t*-test (***, *P* < 0.001). (**G**) Western blot analysis and quantification of Sirt5 protein expression in isolated mitochondrial from HL-1 cells under normal and HR conditions. Two-tailed *t*-test (***, *P* < 0.001).

We next investigated the effects of manipulating Sirt5 expression on mitochondrial homeostasis and cardiac function. The efficiencies of shRNA-mediated knockdown in HL-1 cells, cardiac-specific conditional knockout and adeno-associated virus 9 (AAV9)-mediated cardiac overexpression of *Sirt5* were verified by western blot analyses (fig. S1). In HL-1 cells, MitoTracker staining showed that overexpression of *Sirt5* prevented, whereas knockdown of *Sirt5* exacerbated, mitochondrial fragmentation caused by HR treatment (Fig. 2A). Similarly, mitoSOX labeling indicated that overexpression of *Sirt5* decreased, whereas knockdown of *Sirt5* further increased, the level of mitochondrial superoxide dismutase induced by HR injury (Fig. 2B). Moreover, using ATP assay to evaluate ATP level, JC-1 dye to monitor the formation of mitochondrial J-aggregates, and seahorse technology to measure the mitochondrial oxygen consumption rate (OCR), we found that overexpression of *Sirt5* was able to restore ATP production, increase mitochondrial transmembrane potential, and improve mitochondrial respiration in HR-injured HL-1 cells, whereas knockdown of *Sirt5* had the opposite effects (Fig. 2, C to E). To examine the in vivo function of Sirt5 in myocardial IR injury, C57BL/6 mice with AAV9-mediated cardiac overexpression and cardiac-specific conditional knockout of *Sirt5* were subjected to IR as described above. As revealed by electrocardiography, triphenyltetrazolium chloride (TTC) staining and hematoxylin-eosin (H&E) staining, Overexpression of *Sirt5* significantly improved heart function by increasing ejection fraction and fractional shortening, decreasing infarct size, and reducing inflammatory cell infiltration, whereas knockout of *Sirt5* further aggravated IR-injured cardiac dysfunction and damage (Fig. 2, F to H). Analyses by transmission electronic microscopy (TEM) indicated that overexpression of *Sirt5* prevented IR-disrupted mitochondrial integrity in the myocardium, whereas conditional knockout of *Sirt5* enhanced mitochondrial defects (fig. S2). Together, these results demonstrate that Sirt5 maintains mitochondrial function and exerts a cardioprotective role after IR injury.

**Fig. 2.**
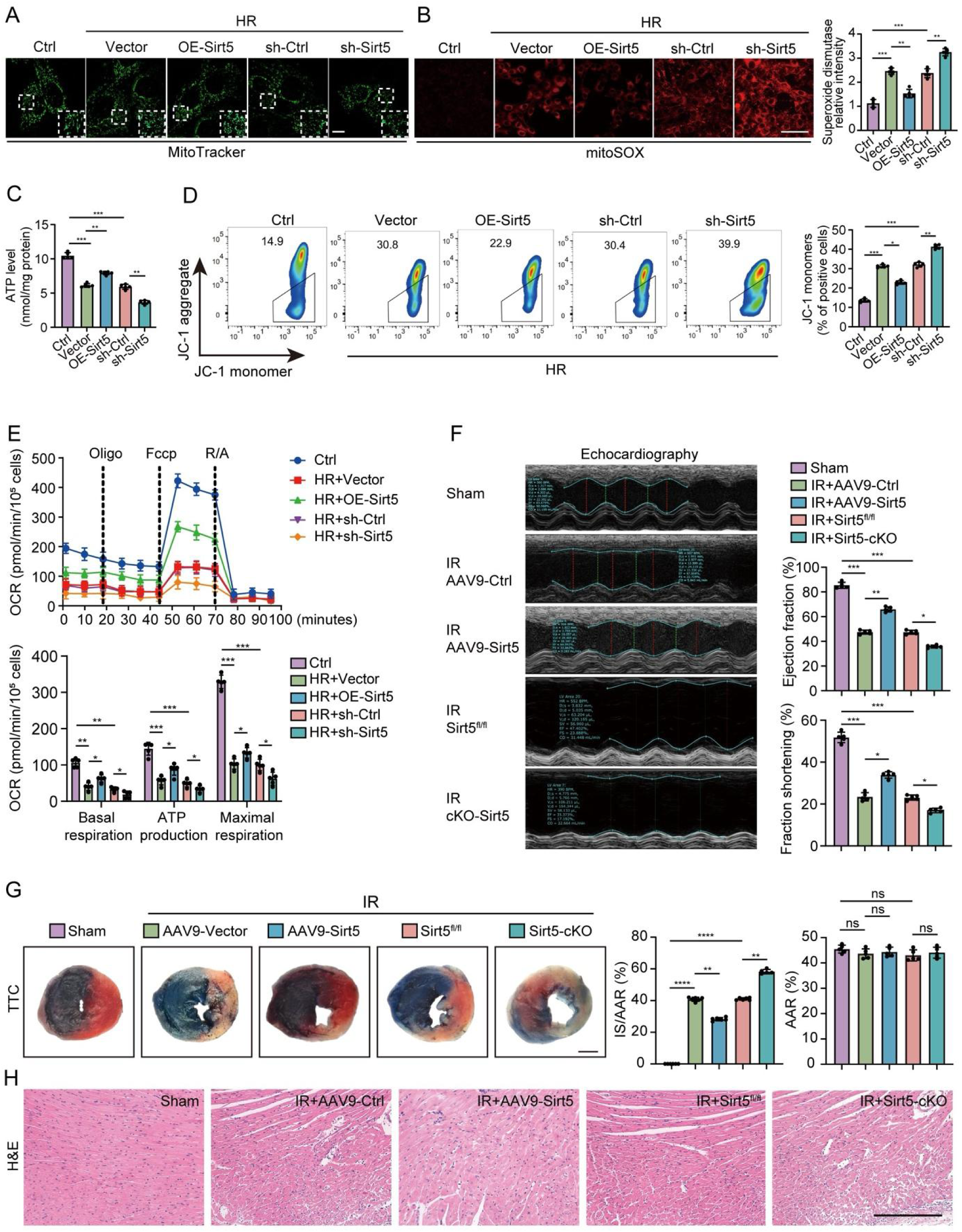
Sirt5 restores mitochondrial and cardiac functions after myocardial IR injury. (**A**) MitoTracker staining of mitochondrial fragmentation in HL-1 cells with *Sirt5* overexpression (OE-Sirt5) or knockdown (sh-Sirt5) for 48 hours followed by HR injury. Insets are enlarged areas. Scale bar, 10 µm. (**B**) Representative images of mitoSOX staining and quantification of mitochondrial ROS in indicated conditions. One-way ANOVA (**, *P* < 0.01; ***, *P* < 0.001). Scale bar, 40 µm. (**C**) Overexpression of *Sirt5* rescues ATP concentration in HL-1 cells subjected to HR injury, whereas knockdown of *Sirt5* further decreases ATP production. One-way ANOVA (**, *P* < 0.01, ***, *P* < 0.01). (**D**) Overexpression of *Sirt5* increases mitochondrial potential in HR-injured HL-cells, as reflected by the reduced formation of JC-1 monomers, whereas knockdown of *Sirt5* shows the opposite effect. One-way ANOVA (**, *P* < 0.01, ***, *P* < 0.01). (**E**) Measurement of the mitochondrial OCR and the quantification of basal respiration, ATP production, and maximal respiration in indicated conditions. One-way ANOVA (**, *P* < 0.01, ***, *P* < 0.01). (**F**) M-mode echocardiograms and quantification of parameters for cardiac function under different conditions in mice 48 hours after IR injury. One-way ANOVA (*, *P* < 0.05; **, *P* < 0.01; ***, *P* < 0.001). (**G**) Evans blue-TTC staining 24 hours after IR injury and quantification show that overexpression of *Sirt5* reduces, whereas conditional knockout of *Sirt5* aggravates, IR-induced myocardial infarct area (IS) relative to myocardial area at risk (AAR). Normal myocardium is stained blue, while ischemic myocardium shows red color. One-way ANOVA (*, *P* < 0.05; **, *P* < 0.01; ***, *P* < 0.001). Scale bar, 1 mm. (**H**) H&E staining compares inflammatory cell infiltration myocardial tissues under different conditions 24 hours after IR injury. Scale bar, 200 µm.

### IR injury increases protein lactylation modifications to disrupt mitochondrial homeostasis

Because sirtuin proteins are important regulators of PTMs, we first set to analyze protein lactylation, crotonylation, succinylation, hydroxybutyrylation and acetylation in IR-injured mouse heart. The results showed that lactylation exhibited a strongest up-regulation (fig. S3, A to E). In line with this observation, higher levels of lactate were present in IR-injured heart and HR-treated HL-1 cells (fig. S4, A and B). The specificity of IR- and HR-induced protein lactylation modifications was confirmed by treating HL-1 cells with dichloroacetate (DCA) and sodium lactate (NaLa), which decreases lactate production and promotes protein lactylation, respectively (*29*, *36*). It has been shown that lysine lactylation represents a dominant and a novel type of PTM driven by lactate (*29*). Consistently, analyses by western blot, immunofluorescence and immunohistochemistry using an antibody against pan-lysine lactylation (pan-Kla) indicated that DCA inhibited, whereas NaLa enhanced, IR- and HR-induced protein pan-Kla modifications (Fig. 3, A to D). In myocardial tissues subjected to IR injury, we also detected a decreased or an increased lactate concentration following treatment with DCA or NaLa, respectively (Fig. 3E). Interestingly, overexpression of *Sirt5* reduced the extent of protein pan-Kla in HR-treated HL-1 cells or IR-injured myocardial tissues, whereas knockdown or knockout of *Sirt5* produced the opposite effects (Fig. 3, F to I). Therefore, we can conclude that the decreased expression of Sirt5 in injured myocardial tissues leads to increased protein lysine lactylation modifications, which may impair mitochondrial activity and damage heart function.

**Fig. 3.**
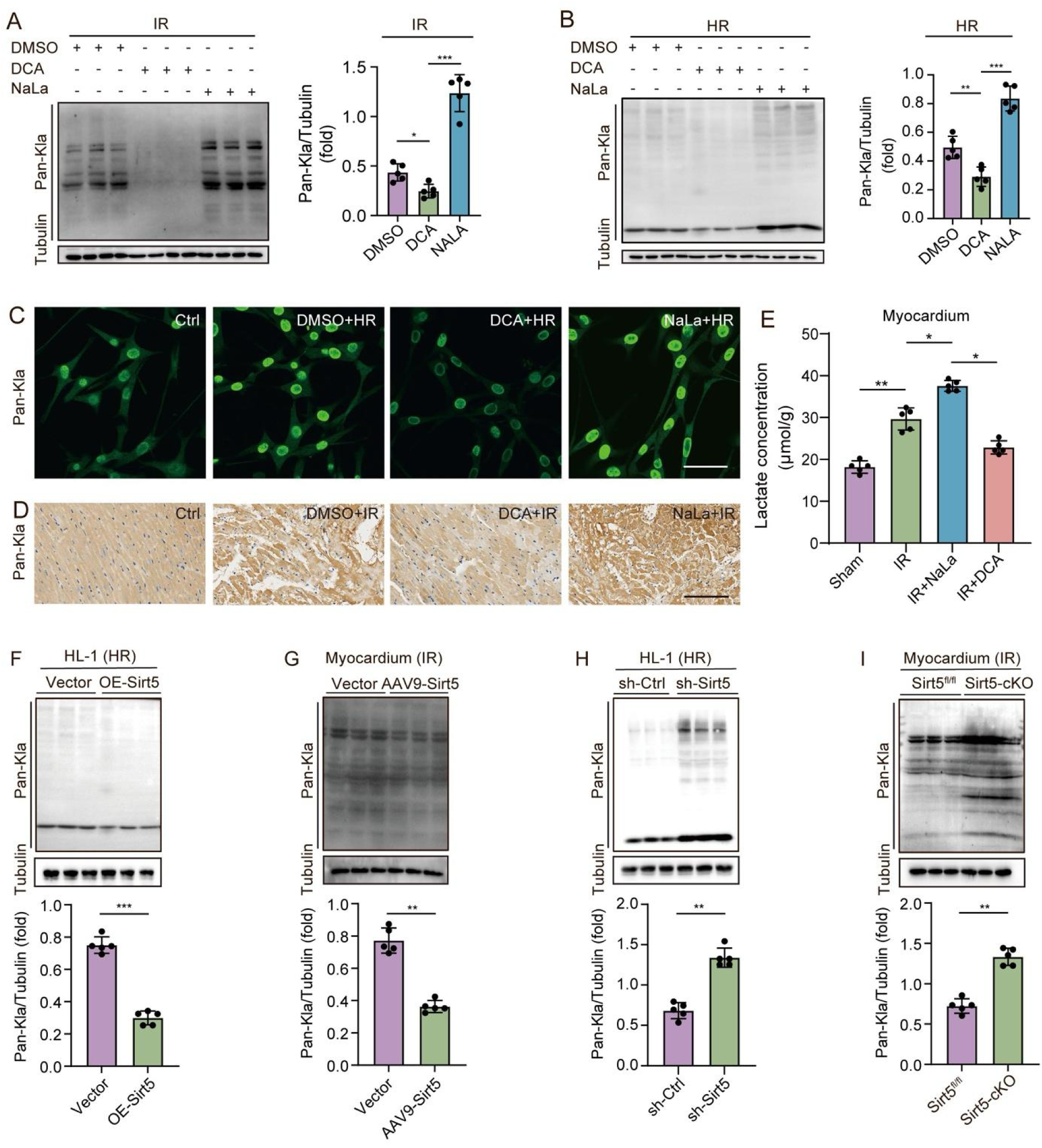
Sirt5 inhibits protein lactylation modifications under HR and IR conditions. (**A**) Analysis of pan-Kla in myocardial tissues from sham and IR-injured mice previously treated with DCA (100 mg/kg) for 15 days or NaLa (0.2 g/kg) for 14 days. One-way ANOVA (*, *P* < 0.05; ***, *P* < 0.001). (**B**) Analysis of pan-Kla in control and HR-injured HL-1 cells previously treated with DCA (20 mM) or NaLa (20 mM) for 24 hours. One-way ANOVA (**, *P* < 0.01; ***, *P* < 0.001). (**C**) Immunofluorescence analysis of pan-Kla in HR-injured HL-1 cells previously treated with DCA or NaLa for 24 hours. Scale bar: 40 µm. (**D**) Immunohistochemistry analysis of pan-Kla in myocardial tissues from sham and IR-injured mice previously treated with DCA or NaLa. Scale bar: 200 µm. (**E**) Quantification of lactate levels in IR-injured mouse myocardial tissues previously treated with NaLa or DCA. One-way ANOVA (*, *P* < 0.05; ***, *P* < 0.001). (**F** to **I**). Overexpression of *Sirt5* inhibits, whereas knockdown or knockout of *Sirt5* increases, protein pan-Kla in HR-treated HL-1 cells and IR-injured myocardial tissues. The analyses were performed 48 hours after transfection in HL-1 cells, 14 days after tail vein injection of AAV9-Sirt5 or 5 days after tamoxifen administration. Two-tailed *t*-test (**, *P* < 0.01, ***, *P* < 0.001).

We thus analyzed the effects of protein lactylation changes on mitochondrial homeostasis. As expected, lowering the lactate concentration in HR-treated HL-1 cells by DCA alleviated the severity of mitochondrial fragmentation, decreased the level of mitochondrial superoxide dismutase, ameliorated ATP production, and increased mitochondrial transmembrane potential. Conversely, increasing protein lactylation modifications by NaLa treatment further deteriorated mitochondrial homeostasis and function (fig. S5, A to D). Analyses by TEM of heart sections from mice subjected to IR injury showed that intraperitoneal injection of DCA improved the structural organization of mitochondria, whereas NaLa further disrupted mitochondrial integrity (fig. S5E). In addition, DCA significantly reduced infarct size and infiltration of inflammatory cells as shown by TTC and histological staining (fig. S5, F and G), and improved cardiac ejection fraction and fractional shortening as revealed by ultrasound examination (fig. S5H).

### Identification of Sirt5-regulated protein lactylation modifications

We used 4D label-free proteomics technology to identify differentially lactylated proteins in mouse ventricular myocardial tissues following IR injury and Sirt5-interacting proteins in HL-1 cells (Fig. 4, A and B). Heat map analysis indicated differential expression of proteins between sham and IR injury groups (fold change > 1.2 or < 0.67, *P* value < 0.05), with 481 proteins down-regulated and 182 proteins up-regulated (Fig. 4C). Volcano plot analysis showed that there were 191 differentially lactylated proteins between sham and IR-injured groups (fold change > 1.2 or < 0.83, *P* < 0.05), among which 30 lactylated proteins with 37 lactylation sites were up-regulated, whereas 161 lactylated proteins with 429 lactylation sites were down-regulated (Fig. 4D). By gene ontology (GO) analysis, we found that differentially lacylated proteins were enriched in various biological processes, such as the generation of precursor metabolites and energy (Fig. 4E). Functional classification in Clusters of Orthologous Groups (COG) of proteins suggested that they were mostly involved in energy production and conversion (Fig. 4F). It is of note that 40% of up-regulated and 52% of down-regulated lacylated proteins in the IR-injured condition were predicted to be localized in the mitochondria (Fig. 4G).

**Fig. 4.**
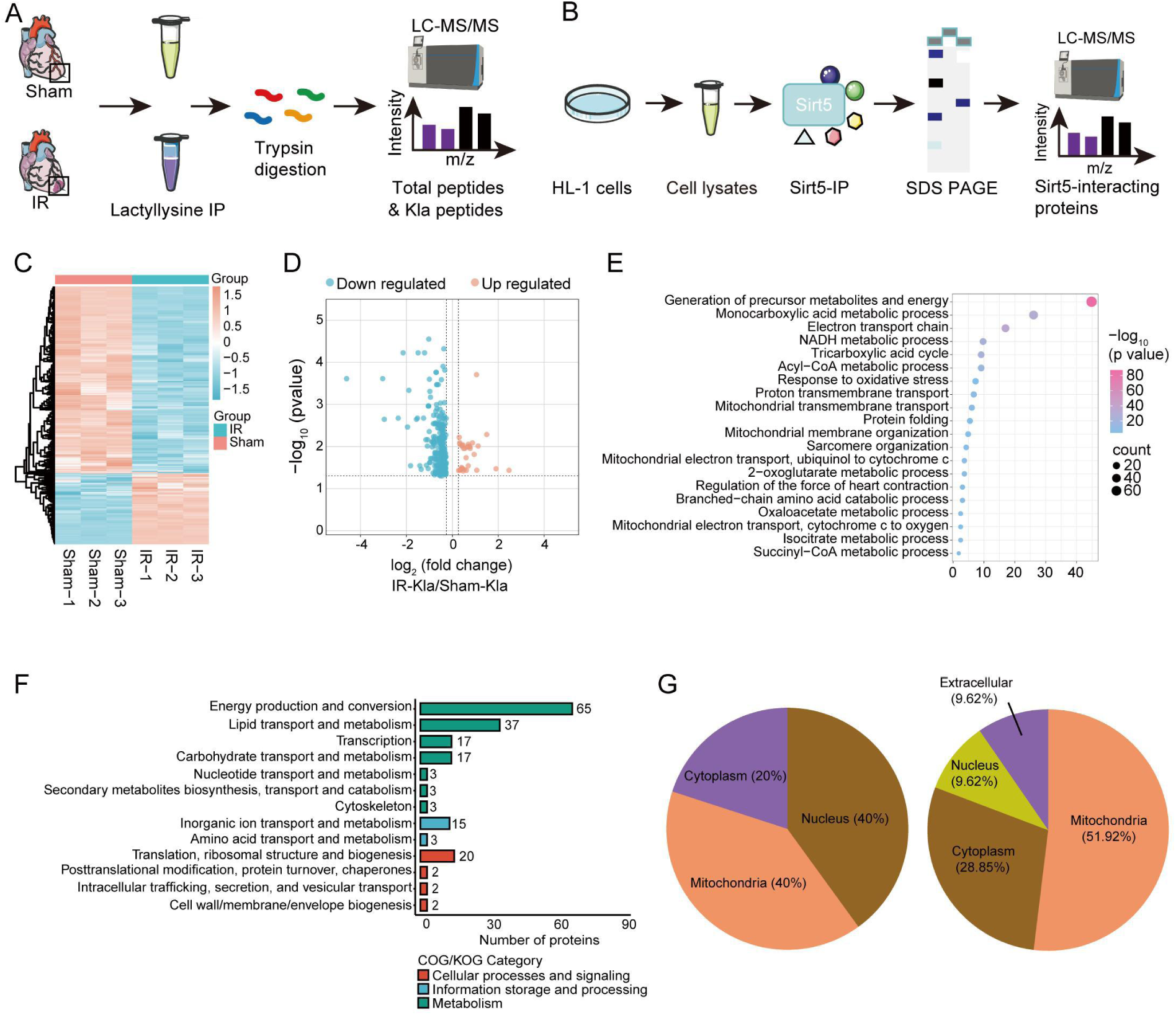
Differential protein lactylation modifications after myocardial IR injury and identification of Sirt5-interacting partners. (**A** and **B**) Flow chart of LC-MS/MS analysis for the identification of protein lactylation modifications in the myocardium and Sirt5-interacting proteins in HL-1 cells. (**C**) Heatmap representation of protein expression changes in mouse myocardial tissues under sham and IR conditions. (**D**) Volcano plot displays differentially lactylated proteins in mouse myocardial tissues under IR condition. (**E**) GO analysis shows the enrichment of differentially lactylated proteins in different biological processes. (**F**) COG category of differentially lactylated proteins. (**G**) Pie charts illustrate the subcellular distribution of differentially lactylated proteins.

Mass spectrometry analysis showed that 92 proteins were co-precipitated with Sirt5. Interestingly, among all potential Sirt5-interacting proteins, only the inner mitochondrial protein ANT2 exhibited an up-regulated lactylation modification in the IR-injured cardiac tissue (Fig. 5A), thus correlating decreased Sirt5 expression with increased ANT2 lactylation in IR-injury. Collision-induced dissociation (CID) analysis indicated that the lysine 163 (K163) residue of ANT2 within a conserved GLGDCLV(Kla)IYK motif represents a major site of lactylation modification (Fig. 5, B and C). We then used the HDOCK software for protein-protein docking and the protein structure information from the Research Collaboratory for Structural Bioinformatics (RCSB) database to predict the mode of interaction between Sirt1-7 and ANT2. Although all Sirt proteins were able to interact with ANT2 (Fig. 5D, fig. S6), Sirt5 showed the strongest binding strength, displaying a more negative docking score compared to the other Sirt proteins, suggesting a strongest interaction (Fig. 5, D and E). Further analysis indicated that ANT2 can form hydrogen bond and salt bridge with Sirt proteins, especially with Sirt5, suggesting a stable interaction. Co-immunoprecipitation (Co-IP) experiments further confirmed the physical interaction between endogenous Sirt5 and ANT2 proteins in HL-1 cells and myocardial tissues (Fig. 5, F and G), as well as between HA-tagged Sirt5 and FLAG-tagged ANT2 expressed in HEK293T cells (Fig. 5, H and I). Thus, ANT2 represents a novel interacting partner of Sirt5, consistent with their co-localization in the mitochondrial matrix.

**Fig. 5.**
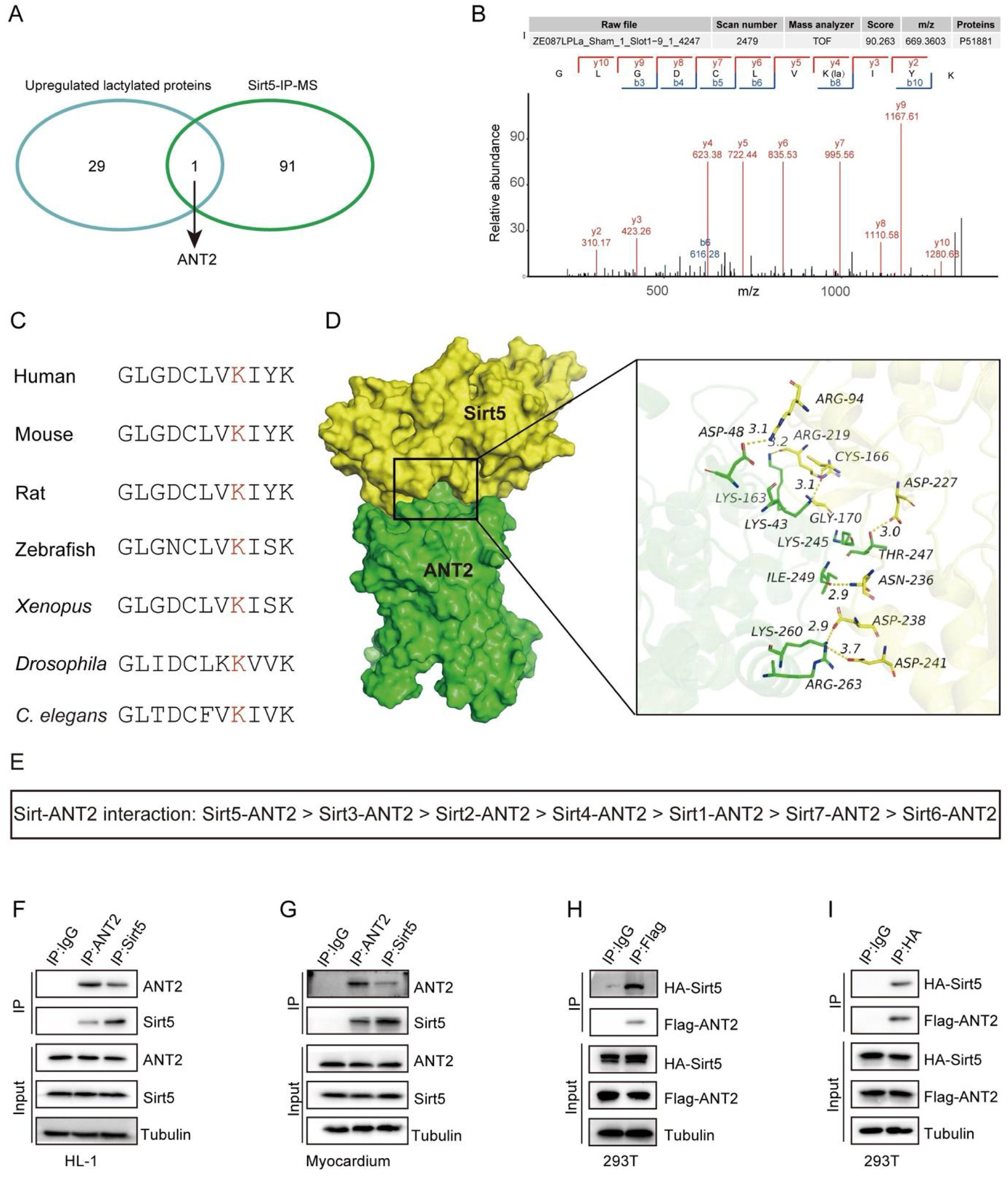
Interaction between Sirt5 and ANT2. (**A**) Venn diagram shows the intersection of up-regulated lactylated proteins and Sirt5-interacting proteins. (**B**) CID analysis of lysine lactylation modification sites in ANT2. (**C**) Conservation of the putative lysine lactylation motif in ANT2 among invertebrate and vertebrate species. (**D**) Docking result shows the protein-protein interaction between ANT2 and Sirt5, with enlarged ribbon structure diagram showing the interaction region on the right. (**E**) Analysis of docking score shows the highest interaction between Sirt5 and ANT2. (**F** and **G**) Co-IP analysis of the interaction between endogenous Sirt5 and ANT2 in HL-1 cells and mouse myocardial tissues. (**H** and **I**) Co-IP analysis shows the interaction between epitope-tagged Sirt5 and ANT2 in HEK293T cells.

### Sirt5 inhibits lysine lactylation modifications of ANT2

The injured-induced ANT2 lysine lactylation of ANT2 was not due to an increase in its overall protein level (Fig. 6, A and B). In addition, there were no obvious changes of ANT2 lysine crotonyllysine (Kcr), succinyllysine (Ksuc), acetylation (Kac) and ß-hydroxybutyryllysine (Kbhb) after HR in HL-1 cells or IR in the myocardium (fig. S7), suggesting that ANT2 was more sensitive to lysine lactylation. Indeed, NaLa increased ANT2 lysine lactylation in vitro and in vivo, whereas DCA had an opposite effect (Fig. 6, C to F). As expected, overexpression of *Sirt5* in HL-1 cells or in the myocardium reduced ANT2 lysine lactylation (Fig. 6, G and H). In addition, the Sirt5 activator (MC3138) inhibited, whereas the Sirt5 inhibitor (nicotinamide) promoted, ANT2 lysine lactylation when they were injected into the myocardium or incubated with HL-1 cells (fig. S8). These results strongly demonstrate that Sirt5 inhibits ANT2 lysine lactylation modification.

**Fig. 6.**
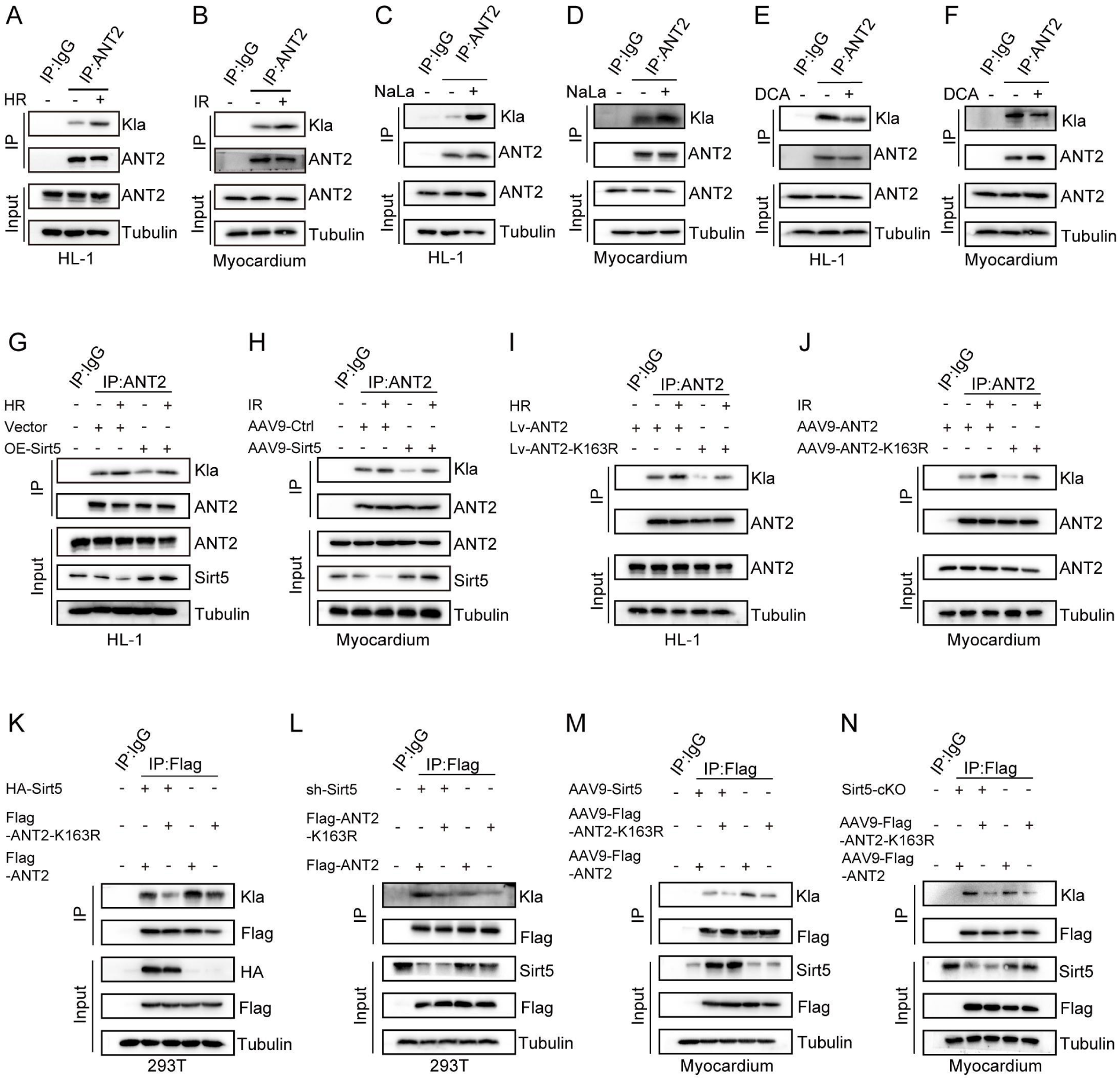
ANT2 is a target of Sirt5-regulated lactylation. (**A** and **B**) Increased ANT2 lysine lactylation following HR injury in HL-1 cells and IR injury in mouse myocardial tissues. (**C** and **D**) Increased ANT2 lysine lactylation 24 hours after NaLa treatment in HL-1 cells and 14 days after NaLa treatment in mouse myocardial tissues. (**E** and **F**) Decreased ANT2 lysine lactylation in DCA-treated HL-1 cells and mouse myocardial tissues. (**G** and **H**) Overexpression of *Sirt5* in HL-1 cells and mouse myocardial tissues inhibits ANT2 lysine modification in the absence and presence of HR or IR injury. Western blot analysis was performed 48 hours or 14 days after overexpression of *Sirt5* in the HL-1 cell line and myocardium, respectively. (**I** and **J**) ANT2-K163R is less sensitive to HR- and IR-induced lactylation than ANT2 in HL-1 cells and mouse myocardial tissues. Lv-ANT2 and Lv-ANT2-K163R are stable HL-1 cell lines generated through lentivirus-mediated gene transfer. (**K** to **N**) Overexpression of *Sirt5* inhibits, whereas knockdown or knockout of *Sirt5* increases, ANT2 and ANT2-K163R lactylation in HEK293T cells and mouse myocardial tissues.

To further evaluate the importance of ANT2-K163 in lactylation, we substituted this lysine residue by an arginine residue (fig. S9A). Expression of the mutated ANT2 protein (ANT2-K163R) in HEK293T cells showed that it was less sensitive to lysine lactylation compared to ANT2 (fig. S10A). Similar results were obtained in mice using AAV9-mediated expression of ANT2 and ANT2-K163R through injection of purified viruses in the tail vein (fig. S9B and fig. S10B). In HL-1 cells treated by HR and myocardium subjected to IR, ANT2-K163R also showed a weak lysine lactylation compared with the wild-type protein either under control or injured condition (Fig. 6, I and J). Therefore, the K163 residue represents a major site that confers ANT2 lactylation. To determine whether this residue is a target of Sirt5-regulated lactylation, we overexpressed FLAG-tagged ANT2 or ANT2-K163R with or without Sirt5 in HEK293T cells and performed immunoprecipitation using the FLAG antibody. Western blot analysis of lactylated protein levels showed that Sirt5 inhibited lysine lactylation of ANT2 and ANT2-K163R (Fig. 6K). Conversely, knockdown of *Sirt5* by shRNA in HEK293T cells increased lysine lactylation of exogenous FLAG-tagged ANT2 but had a very weak effect on ANT2-K163R (Fig. 6L). Similar results were obtained after AAV9-mediated overexpression and conditional knockout of *Sirt5* in the myocardium (Fig. 6, M and N). It is worth mentioning that overexpression of *Sirt5* could further prevent ANT2-K163R lactylation modification. It is possible that other lysine residues in ANT2 may be also potential sites of lactylation, which should be inhibited by Sirt5 because of its strong activity to remove negatively charged lysine modifications (*37*).

### ANT2-K163R improves mitochondrial and cardiac functions

Since Sirt5 regulates lysine lactylation of ANT2, we further assessed its functional interaction with ANT2 and ANT2-K163R in cardiomyocytes. In shRNA-transfected control and *Sirt5*-knockdown HL-1 cells under HR condition, ANT2-K163R was more potent than ANT2 to improve mitochondrial homeostasis by reducing mitochondrial fragmentation and superoxide dismutase level, while increasing mitochondrial ATP production, transmembrane potential and OCR (Fig. 7, A to D). Similarly, in control (*Sirt5*^fl/fl^) and *Sirt5* conditional knockout mice subjected to IR injury, ANT2-K163R more efficiently improved cardiac function when compared with ANT2. Indeed, results from echocardiography, TTC staining and histological examination indicated that AAV9-mediated overexpression of ANT2-K163R in the myocardium significantly increased ejection fraction and fractional shortening of the heart (Fig. 7E), reduced infarct size (IS) relative to the myocardial area at risk (AAR) but without changes in AAR (Fig. 7F), and decreased inflammatory cell infiltration (Fig. 7G). These beneficial effects of ANT2-K163R were correlated with an improved mitochondrial integrity (fig. S11). Therefore, we can postulate that ANT2-K163R may compete with lactylated ANT2 to protect mitochondrial function in cardiomyocytes. As a result, it can counteract the IR injury-induced increase of ANT2 lysine lactylation, thereby maintaining mitochondrial homeostasis and improving cardiac function in a similar manner as overexpression of Sirt5.

**Fig. 7.**
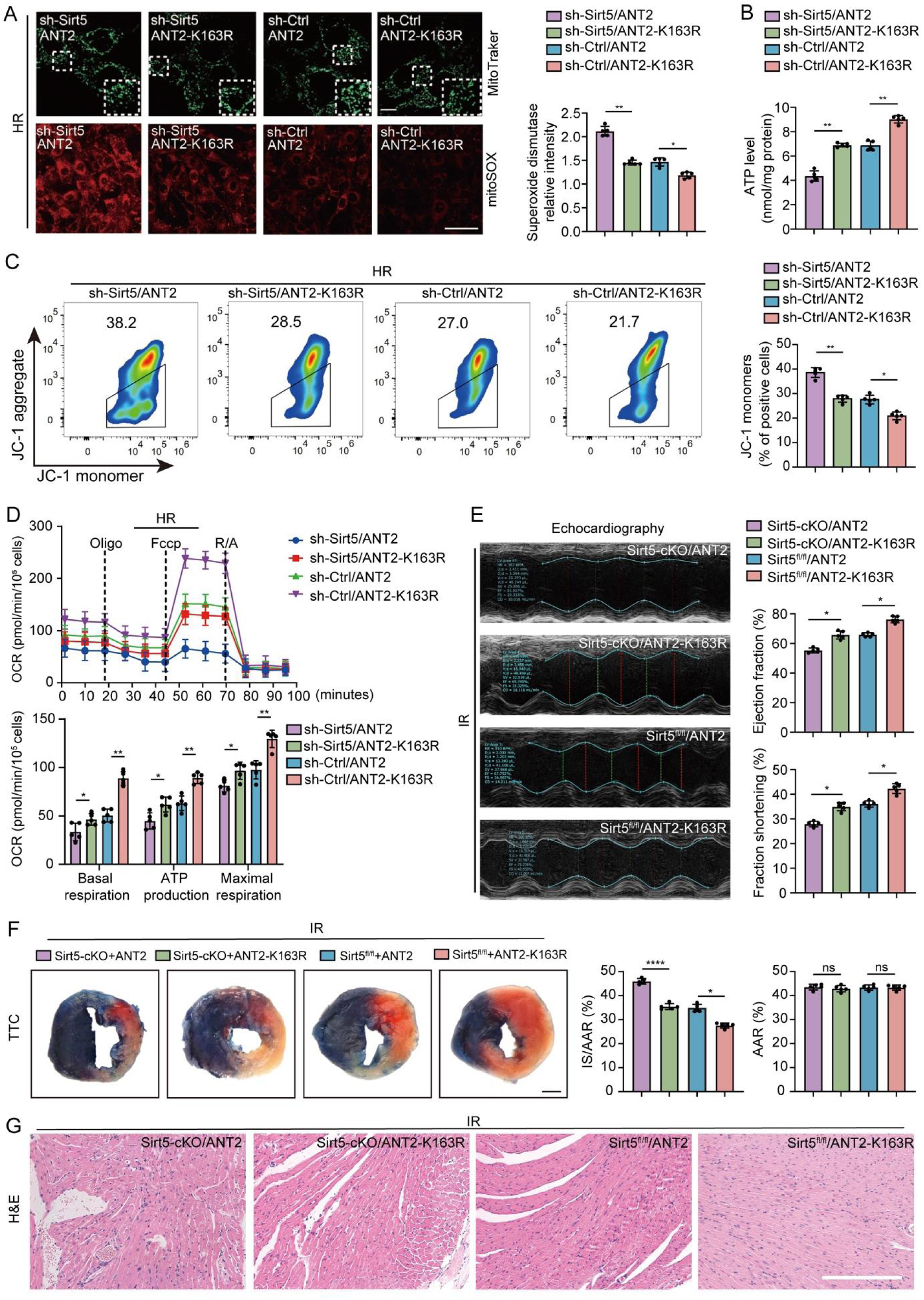
ANT2-K163R improves mitochondrial homeostasis and cardiac function disrupted by Sirt5 deficiency. (**A**) MitoTracker staining and mitoSOX labeling show that ANT2-K163R displays stronger activity to protect mitochondrial integrity and inhibit the increase of ROS level in HR-treated HL-1 cells with or without *Sirt5* knockdown. Quantification of mitochondrial superoxide dismutase intensity is shown on the left. One-way ANOVA (*, *P* < 0.05; **, *P* < 0.01). Scale bars, 5 µm (MitoTracker) and 40 µm (mitoSOX). (**B**) Measurement of ATP levels in HL-1 cells under indicated conditions. One-way ANOVA (**, *P* < 0.01). (**C**) ANT2-K163R is more potent than ANT2 in restoring mitochondrial membrane potential by reducing the formation of JC-1 monomers in HL-1 cells with or without *Sirt5* knockdown. One-way ANOVA (*, *P* < 0.05; **, *P* < 0.01). (**D**) ANT2-K163R shows higher activity than ANT2 to increase mitochondrial OCR in HL-1 cells with or without *Sirt5* knockdown. One-way ANOVA (*, *P* < 0.05; **, *P* < 0.01). (**E**) Ultrasound quantitative imaging compares the activity of ANT2 and ANT2-K163R to improve cardiac function under indicated conditions. One-way ANOVA (*, *P* < 0.05). (**F**) Evans blue TTC staining of ventricular sections show that ANT2-K163R is more effectively than ANT2 to reduce the proportion of infarct size in IR-injured mouse heart with or without *Sirt5* conditional knockout. One-way ANOVA (*, *P* < 0.05; **, *P* < 0.01; ns, not significant). (**G**). H&E staining compares the activity of ANT2 and ANT2-K163R to reduce inflammatory cell infiltration under indicated conditions 24 hours after IR injury. Scale bar, 200 µm.

### Sirt5 regulates the interaction of ANT2 with VDAC1 in mitochondrial homeostasis

To explore the mechanism underlying ANT2 lactylation in cardiac function, we conducted a protein-protein interaction (PPI) network analysis for ANT2 (also known as SLC25A5 in humans) using the STRING database. The results revealed a highest binding affinity of ANT2 with VDAC1 (Fig. 8A). This is in line with the presence of ANT2 and VDAC1 at the inner and outer mitochondrial membrane, respectively, and with their functioning in tandem for efficient transfer of energy metabolites across mitochondria (*38*). In HL-1 cells stably expressing ANT2 or ANT2-K163R, Co-IP experiments using the ANT2 antibody that recognizes both ANT2 and ANT2-K163R indicated that ANT2-K163R was more potent than ANT2 to interact with endogenous VDAC1 (Fig. 8B). Importantly, HR treatment strongly decreased the interaction of ANT2 with VDAC1, whereas the interaction of ANT2-K163R with VDAC1 was less affected (Fig. 8B). In the myocardium, there was also a strong interaction between ANT2-K163R and VDAC1, which was moderately reduced by IR injury (Fig. 8C). These results not only identified a physical interaction between ANT2 and VDAC1, but also revealed a critical role of ANT2 lysine lactylation in regulating the formation of the ANT2-VDAC1 complex.

**Fig. 8.**
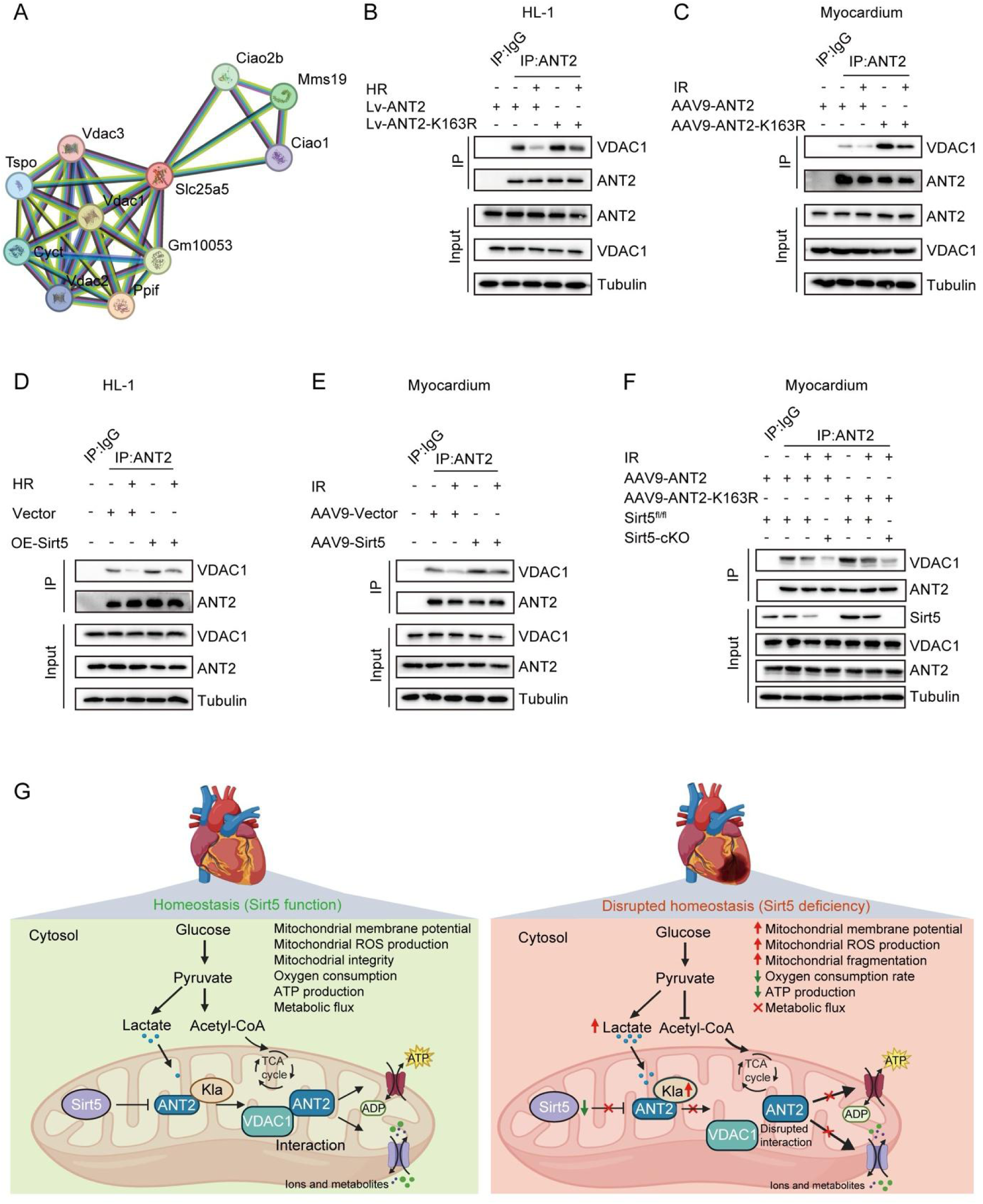
Sirt5-regulated interaction of ANT2 and VDAC1. (**A**) STRING protein network analysis for the identification of ANT2 (SLC25A5) partners. (**B** and **C**) Co-IP shows the interaction of ANT2 and ANT2-K163R with VDAC1 in HL-1 cells and mouse myocardial tissues, which is reduced by HR and IR. The strong interaction of ANT2-K163R with VDAC1 is more resistant to HR and IR. (**D** and **E**) Sirt5 promotes the interactions between endogenous ANT2 with VDAC1 in HL-1 cells and mouse myocardial tissues, which is reduced under HR or IR condition. (**F**) IR injury combined with conditional knockout of *Sirt5* in the myocardium further reduce the interaction between ANT2 and ANT2-K163R with VDAC1. (**G**) Model of Sirt5 function in mitochondrial homeostasis (created with BioRender). Sirt5 regulates ANT2 lysine lactylation and its interaction with VDAC1 to facilitate exchanges of metabolites between the cytosol and mitochondrial matrix. The deficiency of Sirt5 leads to an increased ANT2 lactylation and prevents its interaction with VDAC1, thereby disrupting mitochondrial function and cardiac energy metabolism.

We then determined the role of Sirt5 in the interaction between endogenous ANT2 and VDAC1. In HL-1 cells and myocardial tissues, the Sirt5 activator MC3138 promoted the interaction of ANT2 with VDAC1, but this stimulating effect was affected by HR or IR injury (fig. S12, A and B). Conversely, the Sirt5 inhibitor nicotinamide synergized with HR or IR injury in inhibiting the interaction between ANT2 with VDAC1 (fig. S12, C and D). Consistent with the effects of Sirt5 activator, overexpression of Sirt5 in HL-1 cells or myocardial tissues enhanced the interactions between ANT2 and VDAC1, which was counteracted by HR or IR injury (Fig. 8, D and E). Finally, IR injury combined with conditional knockout of *Sirt5* led to a complete absence of Sirt5 protein and strongly inhibited the interaction of ANT2 or ANT2-K163R with VDAC1 (Fig. 8F). It is well established that ANT2 and VDAC1 are key regulators of mitochondrial functions (*38–40*). There, these results demonstrate that Sirt5 plays an essential role in controlling the interaction between mitochondrial proteins, by modulating lysine lactylation modification of ANT2 (Figure 8G). In addition, there may be a Sirt5-mediated lactylation-independent regulation of ANT2 and VDAC1 interaction. It is possible that the binding of Sirt5 with ANT2 enhances the formation of ANT-VDAC1 complex, which could be weakened by the absence of Sirt5.

## Discussion

In this study we demonstrated that Sirt5 exerts a cardioprotective role of Sirt5 in myocardial IR injury by maintaining mitochondrial homeostasis. We found that myocardial IR injury led to a reduced expression of Sirt5 and mitochondrial dysfunction, which was associated with increased lysine lactylation of ANT2. However, overexpression of Sirt5 was able to improve mitochondrial homeostasis and cardiac function. Mechanistically, Sirt5 binds to ANT2 and inhibits its K163 lactylation, thereby favorizing the interaction between ANT2 and VDAC1 for proper mitochondrial metabolism and function. These findings uncover a regulatory network among key mitochondrial proteins and provide new insights into the molecular mechanism underpinning the maintenance of mitochondrial homeostasis in myocardial IR injury.

Sirtuin proteins are critically involved in protein PTMs essential for cardiac function (*18*, *20*, *21*). Sirt5 is a mitochondrial protein with a very weak deacetylase activity (*25*). Evidence is accumulating that it functions as a major regulator of cellular homeostasis and plays pivotal roles in multiple aspects of myocardial energy metabolism (*37*). We found that Sirt5 exerted a protective role on mitochondrial homeostasis and cardiac function following myocardial IR injury. Our results are consistent with previous studies demonstrating that the loss of Sirt5 causes hypertrophic cardiomyopathy associated with reduced cardiac function and impacts IR injury by increasing the infarct size (*26*, *41*). Indeed, Sirt5 has been shown to preserve cardiac function and mouse survival in response to pressure overload (*42*, *43*). It is also implicated in diverse metabolic pathways related to mitochondrial and cardiac functions, such as glycolysis, fatty acid oxidation, and ROS detoxification (*37*). Sirt5 mostly hydrolizes negatively charged lysine modifications (*22–24*). There are many lines of evidence suggesting that Sirt5 regulates protein lysine succinylation with important contribution to heart metabolism, disease and function (*26*, *27*, *41*, *43*, *44*). However, it is still unclear whether and how Sirt5 is implicated in other types of PTM relevant to the maintenance of mitochondrial homeostasis and cardiac function.

We have provided compelling evidence for a novel role of Sirt5 in protein lysine lactylation, which contributes to maintain mitochondrial homeostasis and protect against IR-induced cardiac injury. First, the loss of Sirt5 in the myocardium leads to increased protein lactylation modifications, which is linked to disrupted mitochondrial and cardiac functions. Second and importantly, Sirt5 physically interacts with ANT2 and inhibits its lactylation modifications mostly ocurring on the K163 residues. Third, overexpression of Sirt5 or inhibition of ANT2 lysine lactylation shows beneficial effects for improving mitochondrial homeostasis and cardiac function. Lactylation has been identified as a novel type of protein PTM and a common histone mark in cells, critically contributing to the metabolic regulation of gene expression and cellular function (*29*). In the neonatal mouse heart, global protein lactylation changes are associated with cardiac metabolic reprogramming and cardiac regeneration (*46*). Histone lactylation can activate glycolysis-related gene expression and reparative transcriptional response to promote cardiac repair and attenuate IR injury (*32*, *47*). However, increased lactate concentrations after myocardial infarction can also induce cardiac fibrosis and exacerbating heart dysfunction by promoting the lactylation modification of proteins associated with endothelial-to-mesynchymal transition (*33*). Thus, the impacts of protein lactylation on the cardiometabolic processes may be dependent on the lactylated substrates.

ANT2 is an inner mitochondrial membrane protein, which is encoded by the *SLC25A5* gene on the X chromosome in humans. It plays a key role in maintaining mitochondrial membrane potential with protecting activity against apoptosis (*39*). However, its function in heart homoeostasis remains largely elusive. We demonstrated that IR injury did not change ANT2 protein levels but specifically increased its lysine lactylation modification, which was correlated with the reduced expression of Sirt5. Importantly, we demonstrated a physical interaction of Sirt5 with ANT2 and identified a conserved lysine residue (K163) in ANT2 as a target of Sirt5-regulated lactylation modification. Thus, the inhibition of ANT2 lysine lactylation by up-regulating Sirt5 activity or by mutating the K163 residue (ANT2-K163R) could restore mitochondrial homeostasis and improve cardiac function after IR injury. These findings suggest that ANT2 lysine lactylation, as the loss of Sirt5 function, causes mitochondrial dysfunction, thereby affecting the heart energy metabolism. Consistently, it has been shown that the deficiency of ANT2 in mice causes mitochondrial disorganization and cardiac failure, likely due to disrupted mitochondrial permeability transition pore (*48*). Therefore, the proper function of ANT2 is important for mitochondrial integrity and cardioprotection. Together, these observations provide the functional evidence that Sirt5 coordinates cardiometabolic networks by regulating ANT2 lysine lactylation. They suggest that modulation of ANT2 lactylation should exert a cardioprotective effect in diseased conditions and represents a potential therapeutic strategy for improving cardiac function after IR injury.

What is the consequence of increased ANT2 lactylation in the myocardium? We uncovered that ANT2 physically interacts with VDAC1, a most abundant outer mitochondrial membrane protein that regulates mitochondrial function in health and disease (*38*, *40*). There are three VDAC isoforms (VDAC1, VDAC2 and VDAC3) in humans and mice. They complex and function in tandem with ANT proteins to regulate exchanges of metabolites, ions, and nucleotides between the cytosol and mitochondrial matrix, contributing to modulating mitochondrial function (*49*). Importantly, lysine lactylation of ANT2 prevents its interaction with VDAC1, whereas the lactylation-resistant ANT2-K163R displays stronger binding with VDAC1. Therefore, this study unravels a novel mechanism regulating VDAC1 and ANT2 complex formation in mitochondrial energy metabolic flux. Although the precise role of VDAC1 in IR injury remains unclear, recent evidence suggests that it may be involved in cardioprotection. In H9c2 cardiomyocytes, knockout of VDAC1 leads to oxidative stress-induced apoptosis and enhanced glycolytic stress (*50*). In ischemic hearts, knockdown of VDAC1 prevents the metabolic switch by affecting the ß-oxidation of long-chain fatty acid (*51*). It is likely that the disruption of ANT2-VDAC1 complex due to an increased ANT2 lysine lactylation modification causes the closure of VDAC1, thus suppressing both entry and exit of mitochondrial metabolites (*52*). Accordingly, inhibiting ANT2 lactylation can promote its interaction with VDAC1, thereby improving mitochondrial function and maintaining cellular homeostasis in IR-injured heart. Nevertheless, we cannot exclude the possibility that the docking of ANT2 in Sirt5 may promote its interaction with VDAC1, independently of lactylation inhibition, because the complete absence of Sirt5 also strongly prevents complex formation between ANT2-K163R and VDAC1?

Having established the protective role of Sirt5 in myocardial IR injury, this study should provide a theoretical basis for the design of new therapeutic strategies in the intervention of heart failure. Targeting Sirt5-regualted ANT2 lactylation may be an option for treating IR-induced cardiac injury. In addition, the findings should help identify potential biomarkers for the diagnosis of cardiac dysfunction. Nevertheless, although we have elucidated a novel mechanism underlying Sirt5 function in maintaining mitochondrial homeostasis to improve cardiac function, how Sirt5 regulates lysine lactylation of ANT2 in cardiac injury and the consequence of disrupted interaction of ANT2 with la VDAC1 in mitochondrial metabolic flux await future investigation.

## Materials and Methods

### Ethical statements

Animal experiments were approved by the Ethics Committee of Guangdong Medical University (permit number AHGDMU-LAC-B-202403-0014) and adhered to the principles of the ARRIVE guidelines. The study involving human biopsy was performed by following the ethical principles of the Declaration of Helsinki and was approved by the Institutional Ethics Committee of the Affiliated Hospital of Guangdong Medical University (protocol number PJKT2024-080). Human cardiac tissue samples were collected with written informed consents of each patient.

### Myocardial IR injury in mice

C57BL/6 mice were obtained from SPF Biotechnology (Beijing, China) and housed under standard conditions. Two-month-old mice were randomly assigned to sham and IR groups. After endotracheal intubation, they were anesthetized with 5% isoflurane for 2 minutes, followed by 2% isoflurane until the completion of surgery. Upon opening of the fourth intercostal space, the left coronary artery was ligated with an 8-0 prolene suture to induce myocardial ischemia. The ligature was tied into a slipknot, and the proximal end was trimmed, leaving 3–4 cm of suture outside the chest cavity. After 45 minutes, the ligature was slowly withdrawn to restore myocardial blood flow for 6 hours.

### Conditional knockout mice

Mice with *loxP* sites flanking the exon 3 of *Sirt5* locus were generated by Shanghai Model Organisms Center (Shanghai, China). Mice with cardiac-specific expression of Cre/Esr1 driven by the α-myosin heavy chain 6 (Myh6) promoter were purchased from The Jackson Laboratory. Cardiac-specific conditional knockout (cKO) of *Sirt5* was induced by administering tamoxifen at a dose of 60 mg/kg via daily intraperitoneal injection for 5 consecutive days. Age- and sex-matched 8–12 weeks mice were used for all experiments. Myocardial genomic DNA was used for genotyping and cardiac-specific deletion of *Sirt5* was confirmed by PCR analysis using primers located on both sides of the *lox*P sites (forward: 5’-CCACGCACCCACCCATCAAGT-3’, reverse: 5’-CTGCCTGCAAGCCGTGTAGAGAAG-3’). Myocardial Sirt5 protein levels in *Sirt5*^ff/ff^ and *Sirt5* cKO mice was also examined by western blot.

### Collection of clinical cardiac tissue samples

Excess right atrial tissues were collected from three patients with tricuspid valve insufficiency during valve replacement surgery at the Affiliated Hospital of Guangdong Medical University, which did not lead to any clinical complication. A control biopsy (approximately 15 mg wet weight) was obtained after the initiation of cardiopulmonary bypass, but prior to the induction of cardiac arrest. An IR injury biopsy of the same weight was collected close to the location of the control sample 10 minutes after the release of the aortic cross-clamp. Samples were immediately frozen in liquid nitrogen and stored at -80°C for late use.

### Transfection of plasmids and shRNAs

HEK293T (Chinese Academy of Sciences cell bank) and HL-1 cells (Millipore) were cultured at 37°C in a humidified incubator with a 5% CO_2_ atmosphere. They were transfected with expression plasmids or gene-specific shRNAs (Table S1) using Lipofectamine 3000 or Lipofectamine RNAiMAX (Invitrogen), respectively. Non-targeting shRNA (sh-Ctrl) served as a specificity control. Cells were subjected to appropriate analyses 48 hours post-transfection.

### HR injury in HL-1 cells

HL-1 cells were cultured in complete medium (Gibco) with 10% of fetal calf serum (Gibco) until 85% to 90% confluence. The complete medium was replaced by DMEM containing 4.5 g/L of D-glucose (Gibco), and cells were incubated under 1% O_2_, 5% CO_2_ and 94% N_2_ atmosphere for 12 hours (hypoxia). Reoxygenation was performed by culturing cells in complete medium under 5% CO_2_ atmosphere for 4 hours followed by different analyses.

### AAV9-mediated gene overexpression in the myocardium

AAV9-packaged ANT2 and ANT2-K163R at a concentration of 2 x 10^11^ viral genomes (vg)/mL were suspended in 100 µL of PBS. Purified viruses were injected into the tail vein using a 29G insulin injection needle, and mice were allowed to recover their body temperature in a warm environment. Successful targeting of myocardial region was verified using AAV9-packaged GFP. Cardiac tissues were subjected to IR injury 14 days after virus injection. Histological examination and TTC staining were performed 24 hours after IR injury, and ultrasound quantitative imaging was acquired 48 hours after IR injury.

### Lentivirus packaging and generation of stable HL-1 cell lines

HEK293T cells were transfected with 3 μg of plasmid carrying the gene of interest, 3 μg of pMD2.D lentivirus envelop plasmid, and 6 μg of PsPax lentiviral packaging plasmid, using Lipofectamine 2000 (Invitrogen). They were first cultured for 6 hours in OptiMEM I reduced serum medium (Gibco). After replacing with complete medium (Gibco), the virus-containing supernatant was collected at 48 hours and 72 hours post-transfection, filtered, and concentrated using Lenti-X concentrator (TaKaRa). HL-1 cells were cultured in complete medium for 24 hours, then in fresh medium with 25 μL of concentrated lentivirus and 10 ng/mL of polybrene (Sigma-Aldrich) for 48 hours. Stable cell lines were established by selection with 4 mg/mL of puromycin (InvivoGen) for 2 weeks and maintained in 1 mg/mL of puromycin.

### Western blot and Co-IP

Tissues and cells were lysed in RIPA buffer added with 1 mM of phenylmethylsulfonyl fluoride (PMSF) as described (*53*). After determining protein concentrations using the bicinchoninic acid (BCA) protein assay kit (Beyotime), the lysates were mixed with loading buffer and boiled at 100°C for 10 minutes. Proteins were separated by 10% or 12% SDS-PAGE, and transferred onto PVDF membranes (Millipore). The membranes were blocked with 5% non-fat milk and incubated at 4°C overnight with primary antibodies, followed by peroxidase-conjugated secondary antibodies. Protein bands were visualized using Luminata Western HRP substrate (Millipore) and Tanon 5200 chemiluminescence imaging system (Shanghai).

For Co-IP, approximately 500 µg of proteins in IP lysis buffer (25 mM Tris-HCl, 150 mM NaCl, 1% Triton X-100, 2 mM EDTA, 1 µg/mL leupeptin, 1 µg/mL aprotinin, 1 mM Na_3_PO_4_, 3 mM Na_4_P_2_O_4_, 5 mM NaF, 1 mM PMSF, pH 7.5) were mixed with 2 µg of an appropriate antibody (Table S2) at 4°C overnight. Subsequently, 30 µL of protein A/G agarose beads (Thermo Fisher Scientific) were added to the mixture, which was further incubated for 4 hours. The beads were then washed several times with lysis buffer and boiled in 2 x sample buffer. The supernatant was subjected to SDS-PAGE electrophoresis followed by western blot analyses.

### Immunofluorescence and immunohistochemistry

Tissues and cells were fixed in 4% paraformaldehyde and permeabilized with 0.5% Triton-X100 in the presence of 5% normal goat serum (Solarbio). Following overnight incubation with primary antibodies at 4°C, the samples were washed several times with PBS. After incubation with secondary antibodies, nuclei were stained with Hoechst33342 or DAPI (Merck). Images were acquired using a FV3000 confocal microscope (OLYMPUS).

For immunohistochemistry, myocardial tissues were fixed in 4% formaldehyde for 18 hours and embedded in paraffin. Sections were deparaffinized using xylene and rehydrated in decreasing concentrations of ethanol. Antigen retrieval was performed by treating the sections with antigen retrieval solution (Beyotime) for 20 minutes in a steamer. Endogenous peroxidase activity was quenched using 4% H_2_O_2_, followed by blocking with QuickBlock (Beyotime). Sections were incubated with primary antibodies at 4°C overnight, washed several times with PBS containing 0.1% Tween-20, and incubated with peroxidase-conjugated secondary antibodies (Beyotime). Labelled proteins were visualized using 3,3’-diaminobenzidine substrate.

### MitoTacker staining

HL-1 cells were grown on glass coverslips and stained using either 100 or 200 nM of Mito-tracker green (Cell Signaling Technology) in an incubator at 37°C with a 5% CO_2_ atmosphere for 30 minutes, by following the manufacture’s protocol. Images were captured using a confocal laser scanning microscope at a wavelength of 488 nm, and data was processed using the Zen software (Zeiss).

### Flow cytometry

HL-1 cells were washed with PBS and collected after trypsin digestion. Subsequently, 1 mL of JC-1 staining working solution (Thermo Fisher Scientific) was added to collected cells and thoroughly mixed. After incubation at 37°C for 20 minutes, the cells were washed twice with 1 x JC-1 staining buffer. Flow cytometry analysis was performed using a BD FACSCanto II flow cytometer and FlowJo software (Becton, Dickinson and Company).

### Seahorse extracellular flux assay

This was performed using the Seahorse XF Cell Mito Stress Test kit (Agilent Technologies) by following the manufacturer’s protocols. Briefly, HL-1 cells were seeded onto a Seahorse XF 24-well culture microplate (2 x 10^4^ cells per well) and cultured overnight. They were rinsed twice with detection solution and maintained in the XF assay medium. Solutions of 1 mM oligomycin, 2 mM p-trifluoromethoxy carbonyl cyanide phenylhydrazone, and 0.5 mM rotenone plus antimycin A were successively added to the culture at defined time intervals. Mitochondrial OCR was analyzed using the Seahorse XFe 24 Extracellular Flux Analyzer (Seahorse Bioscience), and the results were expressed as pmol/minute as described (*54*).

### Evans blue-TTC staining

Mice were subjected to anesthesia 24 hours post-IR injury, and the left anterior descending coronary artery was temporarily blocked at the site of the initial ligation. Subsequently, 1 mL of 1% Evans blue solution (Sigma-Aldrich) was injected into the left ventricular cavity. The heart was rapidly removed, rinsed three times to eliminate excess dye, flash-frozen, and cut into 1 mm thick slices. The sections were then soaked in 1% TTC solution (Sigma-Aldrich) and imaged using a digital camera. The infarct size and myocardial area at risk were calculated using digital planimetry and analyzed from multiple consecutive sections using the NIH ImageJ software.

### Measurement of lactate levels

This was performed using the Lactate Colorimetric Analysis kit (Biovision), by following the manufacturer’s instructions. Myocardial tissues and HL-1 cells were homogenized in lysis buffer and sonicated on ice at 300 W for 3 minutes (3 seconds on, 7 seconds off). The suspension was centrifuged at 12,000 *g* for 10 minutes at 4°C to collect the supernatant and lactate levels were quantified using a lactate analyzer.

### In-gel digestion and mass spectrometry

The identification of Sirt5-interacting using mass spectrometry were performed as previously described (*55*). HL-1 cells were homogenized in lysis buffer (25 mM Tris-HCl, 150 mM NaCl, pH 7.6, 1% NP-40, 1% sodium deoxycholate, 0.1% SDS) containing a mixture of protease and phosphatase inhibitors, and shaken on ice for 30 minutes. After centrifugation at 13,000 *g* for 15 minutes at 4°C, the cell lysate was subjected to immunoprecipitation through incubation with 2 μg of Sirt5 antibody at 4°C overnight. Protein G-coupled magnetic beads were washed with lysis buffer and incubated with the cell lysate at 4°C for 4 hours. They were then collected by centrifugation, washed five times with the lysis buffer, and boiled with 2 x SDS sample buffer.

Bound proteins were separated by SDS-PAGE and stained with coomassie brilliant blue. Individual lanes were excised and treated with trypsin at 2.5 to 10 µg/µL in 25 mM of NH_4_HCO_3_ at 37°C for 20 hours. Digested peptides were extracted thrice using 60% acetonitrile/0.1% trifluoroacetic acid. The pooled fractions were completely dried using a vacuum centrifuge.

LC-MS/MS analysis was conducted on a Q Exactive mass spectrometer interfaced with an Easy-nLC 1000 liquid chromatography system (Thermo Fisher Scientific). Raw data were searched using the version 2.2 of MASCOT engine (Matrix Science) and compared with the mouse UniProt database using the version 1.4 of Proteome Discoverer (Thermo Fisher Scientific) for protein identification. The peptide mass tolerance and the MS/MS tolerance were set at 20 ppm and 0.1 Da, respectively, and the false discovery rate (FDR) was ≤ 0.01.

### 4D label-free lactylation modification proteomics

Differentially lactylated proteins following myocardial IR injury in mice were identified as previously reported (*56*, *57*). Samples were collected from the left ventricular apex and cells were pulverized in liquid nitrogen. The powder was solubilized in lysis buffer (50 µM PR-619, 1% Triton X-100, 50 µM NAM, 10 µM dithiothreitol, 1% protease inhibitor cocktail, 3 µM trichostatin A, 2 µM EDTA). This mixture was then subjected to high-intensity ultrasonication on ice using an ultrasonic processor (Scientz). After adding an equal volume of saturated phenol-trimethylamine followed by vortexing for 5 minutes, the sample was centrifuged at 5000 *g* for 10 minutes at 4°C. The phenol phase was carefully transferred to a new tube for protein precipitation using saturated ammonium sulfate and methanol at -20°C for at least 6 hours. After centrifugation, the pellet was washed with cold methanol and acetone. Precipitated proteins were dissolved in 8 M of urea, and the concentration was determined using the BCA kit (Beyotime). The solution was subjected to reduction in 5 mM dithiothreitol at 56°C for 30 minutes, and then alkylation in 11 mM iodoacetamide for 15 minutes at room temperature in the dark. The protein sample was treated in 100 mM triethylamine bicarbonate and digested overnight with trypsin at a 1:50 mass ratio. For the enrichment of lactylation-modified peptides, tryptic peptides were dissolved in NETN buffer (100 mM NaCl, 1 mM EDTA, 50 mM Tris-HCl, pH 8.0, 0.5% NP-40) and incubated with anti-lactyllysine antibody conjugated agarose beads (PTM BIO) at 4°C overnight. After thorough washing with NETN buffer and water, peptides were eluted with 0.1% trifluoroacetic acid, vacuum-dried, and analyzed by LC-MS/MS. The spectra were searched against the Mus_musculus_10090_SP_20220107.fasta database and a reverse decoy database using the MaxQuant software. Trypsin/P was set as the enzyme, allowing up to two missed cleavages. The precursor mass tolerance was 20 ppm and the fragment ion tolerance was 0.04 Da. Fixed modification was set for carbamidomethylation on cysteine, and variable modifications included lactylation on lysine and oxidation on methionine. The false discovery rate was kept below 1%.

### Molecular docking

Protein structures for mouse Sirt1 (Q923E4), Sirt2 (Q8VDQ8), Sirt3 (Q8R104), Sirt4 (Q8R216), Sirt5 (Q8K2C6), Sirt6 (P59941), Sirt7 (Q8BKJ9) and ANT2 (P51881) were obtained from the RCSB database (https://www.rcsb.org/). They were processed on the Molecular Operating Environment platform (MOE 2019.1) with the force field selection set to Amber 10, including the removal of water and ions, protonation, addition of missing atoms, completion of missing groups, and protein energy minimization. The HDOCK server was used to predict the relevant binding by employing a hybrid docking strategy and setting each protein as a rigid molecular scaffold. The docking score was calculated using the knowledge-based iterative scoring function provided by ITScorePP, with a more negative value suggesting strong binding. Subsequently, the Pymol2.1 software was used for visual analysis of the model.

### TEM

Myocardial tissues were fixed with a mixture of 2% formaldehyde and 2% glutaraldehyde, followed by post-fixation in 1% osmium tetroxide (SPI-CHEM). Samples were dehydrated with increasing ethanol concentrations and then embedded in resin. Ultrathin sections were cut and stained with uranyl acetate, followed by staining with lead citrate. Sections were examined using a JEM-1400 transmission electron microscope (JEOL).

## Statistical analysis

All data were collected from 2-3 independent experiments with multiple samples. Statistical analysis was performed using one-way ANOVA with Bonferroni multiple comparisons test or unpaired two-tailed *t*-test. Statistical significance is indicated in the figures and legends.

## Acknowledgments

We thank the core facility and service platform at the Second Affiliated Hospital and the Affiliated Hospital of Guangdong Medical University for assistance with confocal and TEM imaging and animal care.

## Author Contributions

Conceptualization: SL, LZ

Methodology: SL, SS, YH, KD, SC, JC, CW, YW, GM

Investigation: SL, SS, YH Visualization: SL, SS, YH

Supervision: LZ, DLS, LY

Funding acquisition: LZ

Writing—original draft: SL, DLS

Writing—review & editing: DLS, SL, LZ

## Data availability

All data are included in the article and supporting information. Further information and requests for resources and reagents should be directed to the corresponding authors.

## Supplementary Materials

**Fig. S1.**
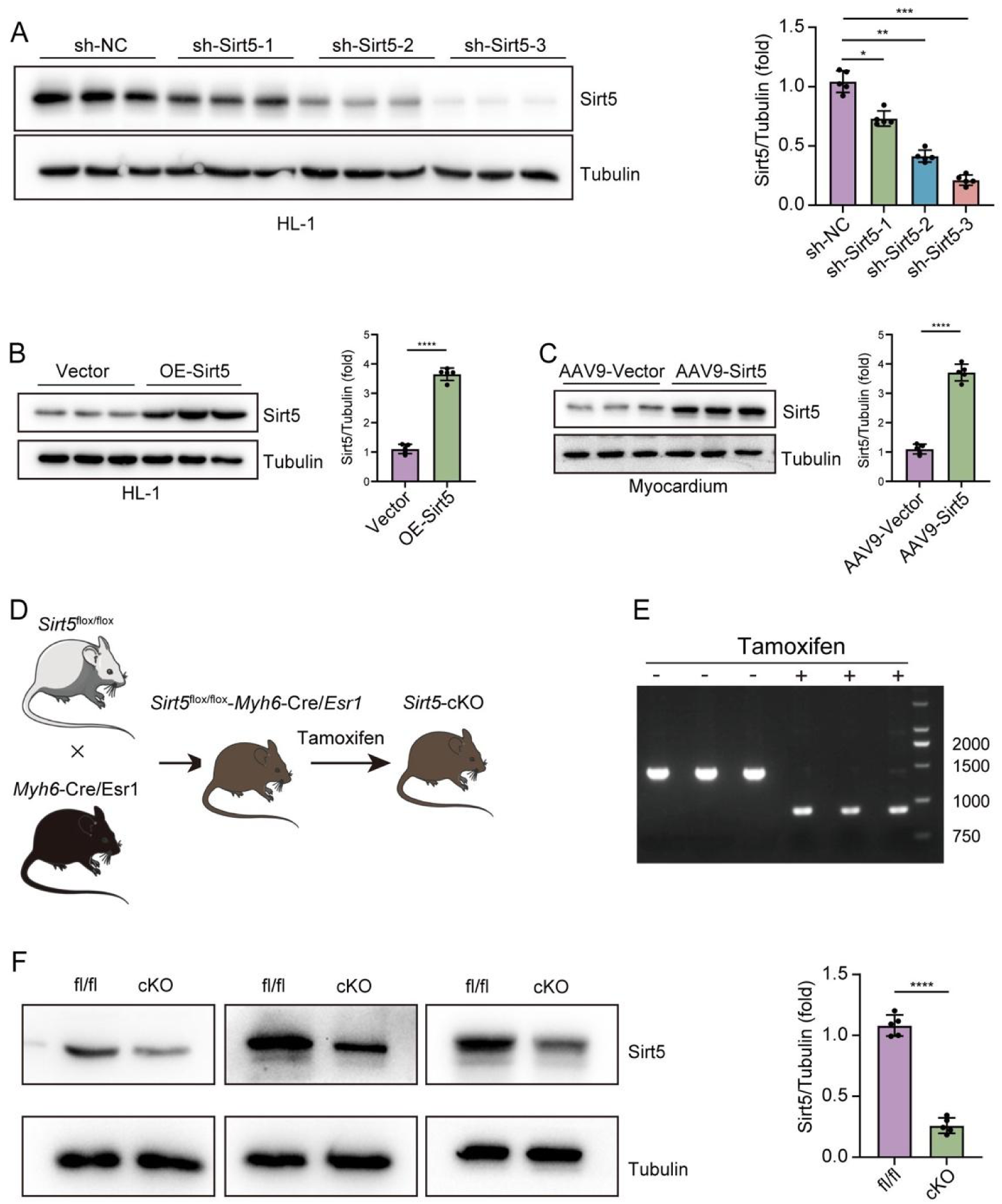
Validation of *Sirt5* overexpression and knockdown or knockout. (**A**) Western blot analysis and quantification shows the efficiency of three shRNAs targeting *Sirt5* in HL-1 cells 48 hours after transfection. The sh-Sirt5-3 was used throughout the study. One-way ANOVA (*, *P* < 0.05; **, *P* < 0.01; ***, *P* < 0.001). (**B**) Western blot analysis and quantification shows increased Sirt5 protein levels after plasmid transfection in HL-1 cells. Two-tailed *t*-test (****, *P* < 0.0001). (**C**) AAV9-mediated overexpression of Sirt5 in the mouse myocardium 14 days after injection of the viruses into the tail vein. Two-tailed *t*-test (****, *P* < 0.0001). (**D**) Procedure for the generation of cardiac-specific *Sirt5* conditional knockout mice. (**E**) PCR-mediated genotyping to verify the deletion of the floxed exon 3 in the *Sirt5* locus, which produces a 1508 bp or 847 fragment before and after administration of tamoxifen, respectively. (**F**) Reduction of Sirt5 protein levels 5 days after tamoxifen treatment in three independent mice. Two-tailed *t*-test (****, *P* < 0.0001).

**Fig. S2.**
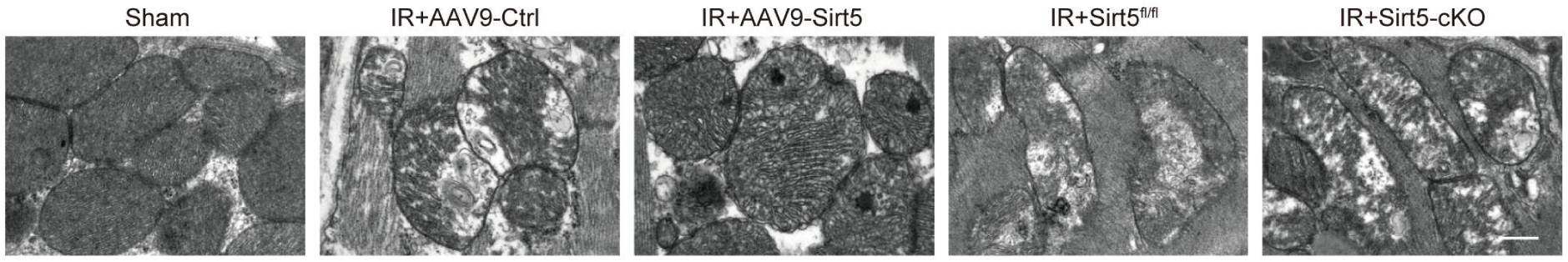
Sirt5 protects mitochondrial integrity under IR condition. IR injury of the myocardium was induced 14 days after injection of AAV9-Sirt5 and 5 days after administration of tamoxifen. Overexpression of *Sirt5* rescues IR-induced mitochondrial disorganization, whereas knockout of *Sirt5* enhances IR-induced mitochondrial defects. Scale bar, 500 nm.

**Fig. S3.**
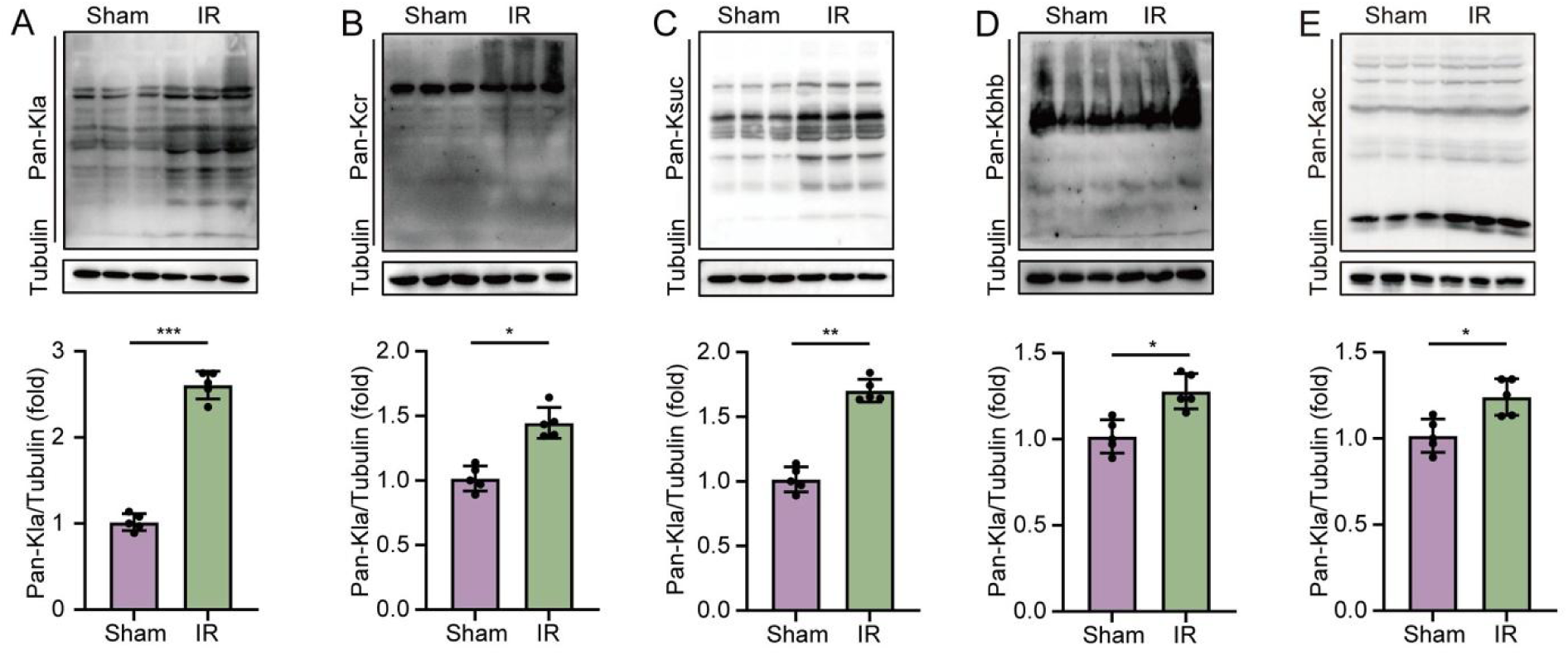
IR-induced PTMs in the myocardium. (**A** to **E**) Western blot analyses of protein lactylation (Kla), crotonyllysine (Kcr), succinyllysine (Ksuc), ß-hydroxybutyryllysine (Kbhb) and acetylation (Kac) modifications in IR-injured mouse myocardial tissues. Two-tailed *t*-test (*, *P* < 0.05; **, *P* < 0.01; ***, *P* < 0.001).

**Fig. S4.**
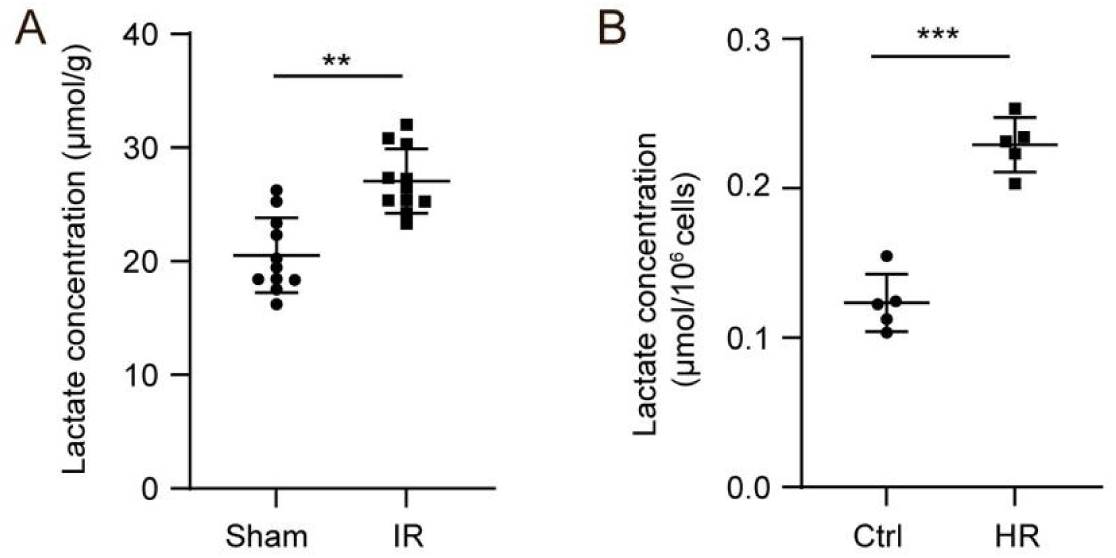
Increase of lactate levels after IR and HR. (**A**) Quantification of lactate levels in IR-injured mouse myocardial tissues. Two-tailed *t*-test (**, *P* < 0.01). (**B**) Quantification of lactate levels in HR-treated HL-1 cells. Two-tailed *t*-test (***, *P* < 0.001).

**Fig. S5.**
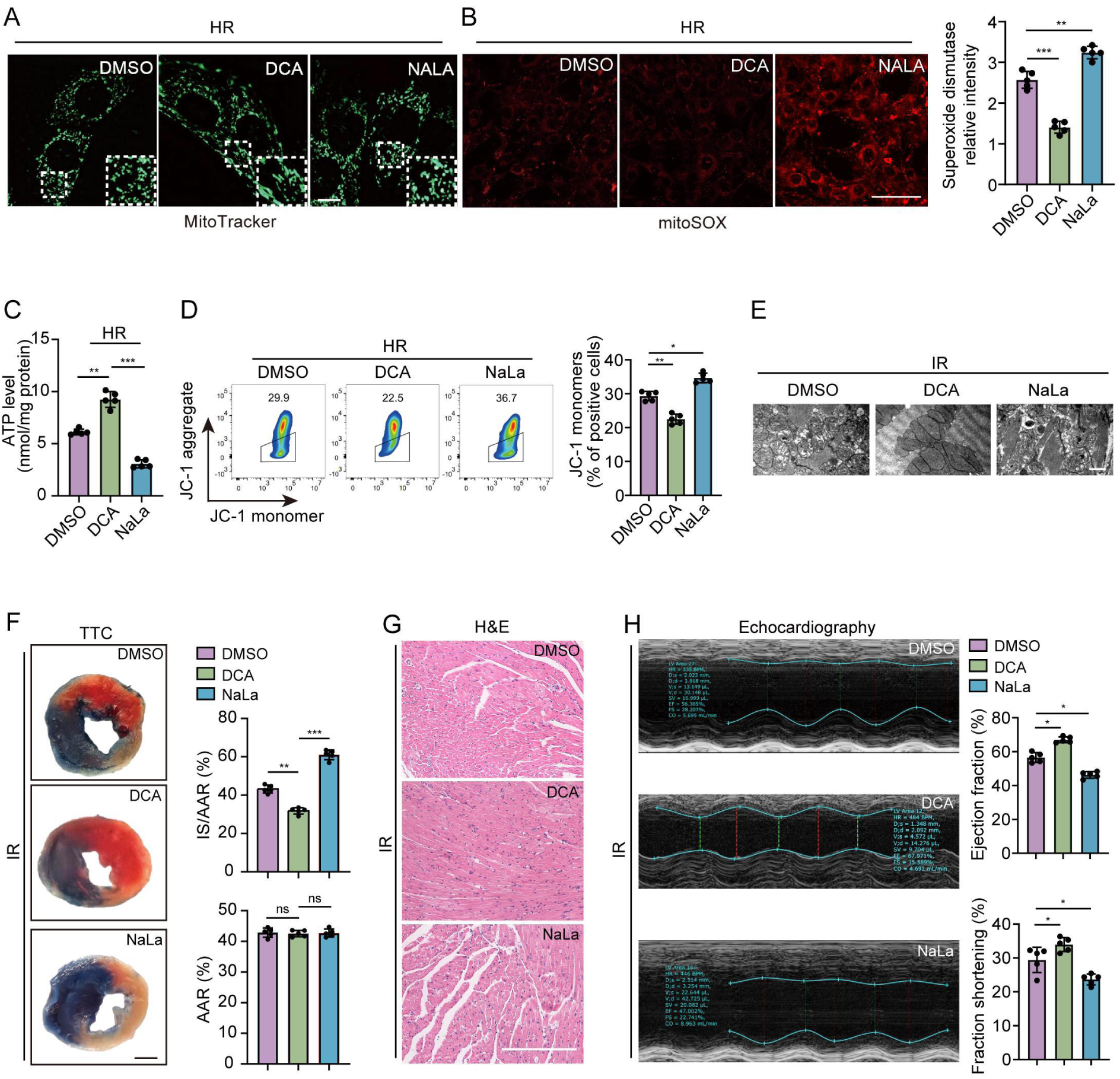
Improvement of mitochondrial and cardiac functions by inhibiting lactate production and lactylation modification. (**A**) DCA reduces, whereas NaLa enhances, the severity of mitochondrial fragmentation in HR-injured HL-1 cells. Scale bar, 10 µm. (**B**) DCA reduces, whereas NaLa enhances, mitochondrial ROS in HR-injured HL-1 cells. One-way ANOVA (**, *P* < 0.01; ***, *P* < 0.001). Scale bar, 40 µm. (**C**) ATP production in HR-injured HL-1 cells treated with DCA or NaLa. One-way ANOVA (**, *P* < 0.01; ***, *P* < 0.001). (**D**) DCA improves mitochondrial membrane potential in HR-injured HL-1 cells by reducing the formation of JC-1 monomers. One-way ANOVA (*, *P* < 0.05; **, *P* < 0.01). (**E**) TEM analyses show that DCA improves, whereas NaLa aggravates, mitochondrial integrity in IR-injured mouse myocardial tissues. Scale bar, 500 nm. (**F**) TTC staining shows that DCA reduces, whereas NaLa further increase, the infarct size (IS) relative to the myocardial area at risk (AAR) from IR-injured mice, but without effect on the AAR. One-way ANOVA (**, *P* < 0.01; ***, *P* < 0.001; ns, not significant). (**G**) DCA reduces, whereas NaLa aggravates, inflammatory cell infiltration in IR-injured mouse myocardial tissues. TTC staining was performed 24 hours after IR injury. Scale bar, 200 µm. (**H**) DCA improves heart function by increasing the percentage of ejection fraction and fraction shortening in IR-injured mice, whereas NaLa shows the opposite effect. Echocardiography was performed 48 hours after IR injury. One-way ANOVA (*, *P* < 0.05).

**Fig. S6.**
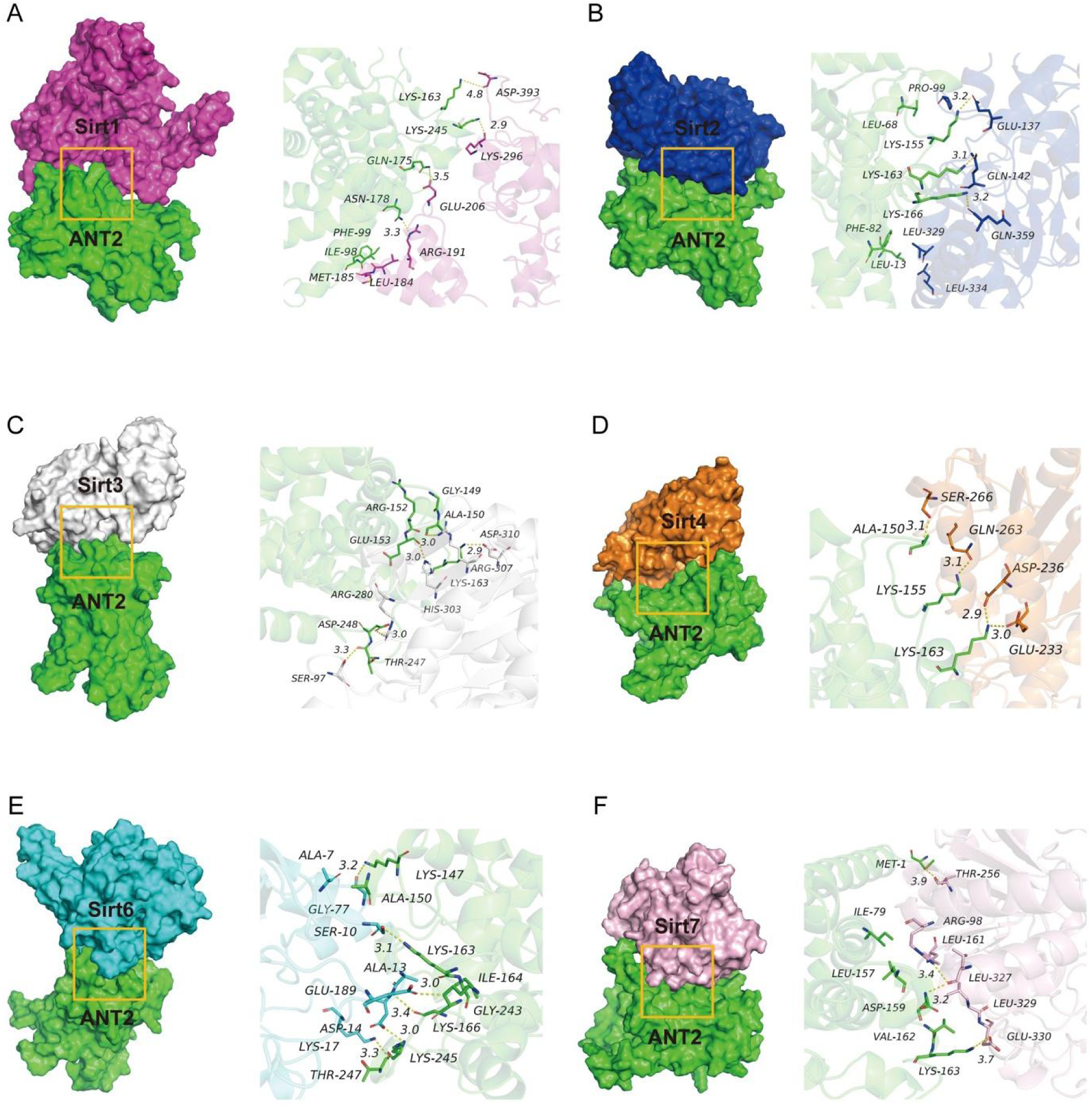
Molecular docking of ANT2 interaction with Sirt proteins. (**A**) Sirt1 interaction with ANT2. (**B**) Sirt2 interaction with ANT2. (**C**) Sirt3 interaction with ANT2. (**D**) Sirt4 interaction with ANT2. (**E**) Sirt6 interaction with ANT2. (**F**) Sirt7 interaction with ANT2. The 3D interactions of ANT2-K163 with Sirt proteins are shown on the right.

**Fig. S7.**
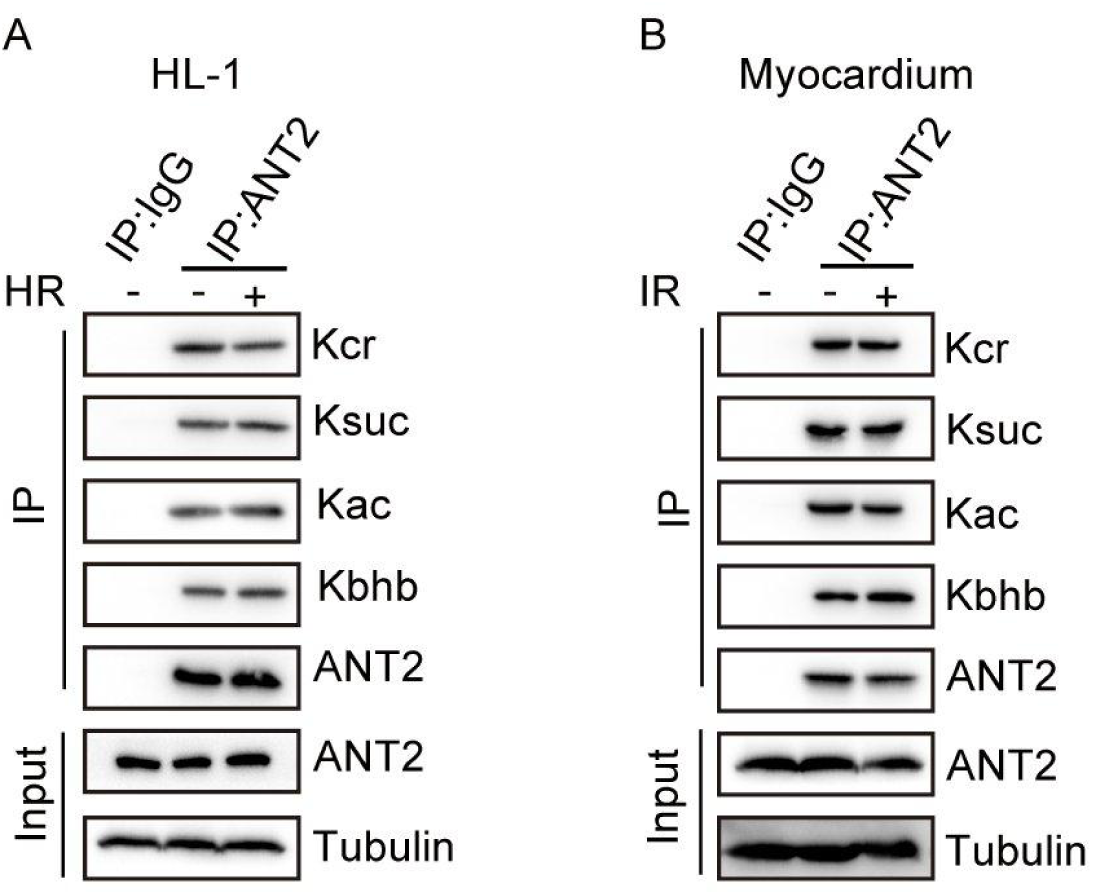
Analysis of ANT2 post-translational modifications. (**A** and **B**) Western blot analyses of ANT2-Kla, ANT2-Kcr, ANT2-Ksuc, ANT2-Kbhb and ANT2-Kac following immunoprecipitation of ANT2 in HL-1 cells and mouse myocardial tissues.

**Fig. S8.**
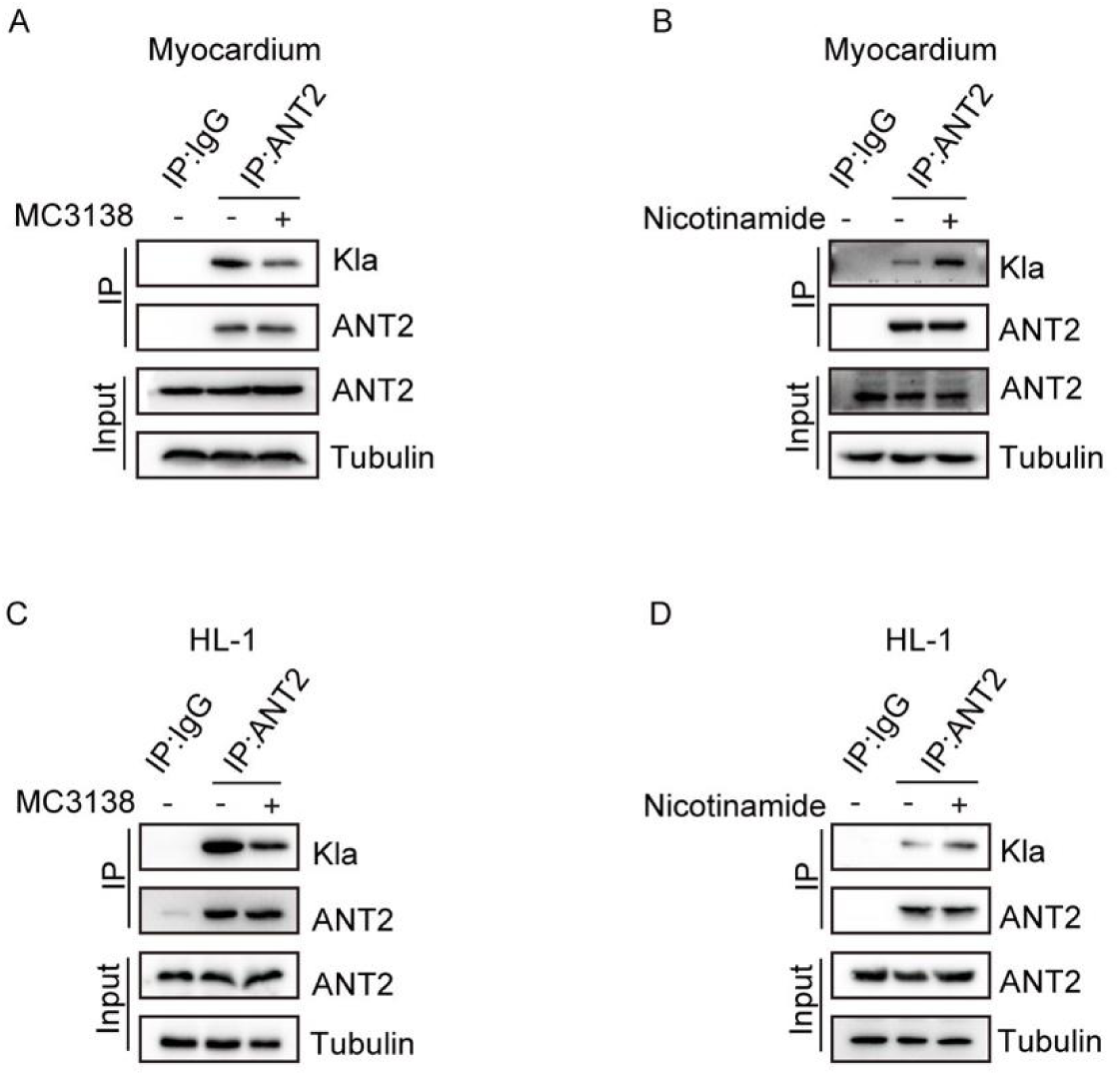
Regulation of ANT2 lysine lactylation by Sirt5 activator and inhibitor. (**A** and **B**) Decreased and increased ANT2 lysine lactylation 14 days following treatment of mouse myocardial tissues with Sirt5 activator MC3138 and inhibitor nicotinamide, respectively. (**C** and **D**) Decreased and increased ANT2 lysine lactylation 24 hours following treatment of HL-1 cells with Sirt5 activator MC3138 and inhibitor nicotinamide, respectively.

**Fig. S9.**
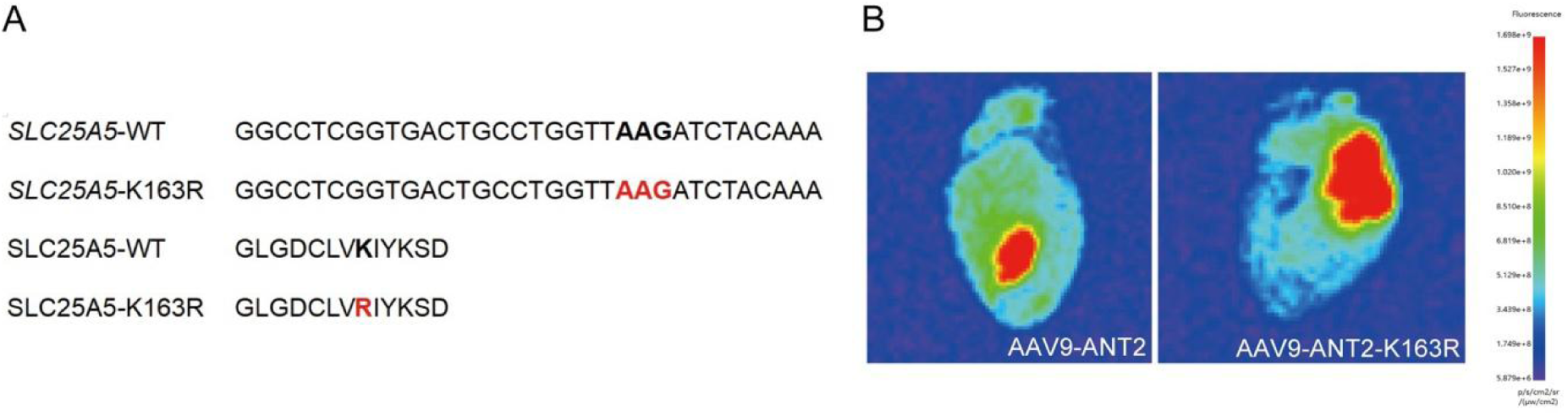
AAV9-mediated cardiac expression of ANT2 and ANT2-K163R. (**A**) Nucleotide and amino acid sequences showing the substitution of the lysine residue in human SLC25A5 (ANT2) by an arginine residue. (**B**) Fluorescence images show the efficiency of AAV9-mediated expression of ANT2 and ANT2-K163R in the heart 14 days after tail vein injection.

**Fig. S10.**
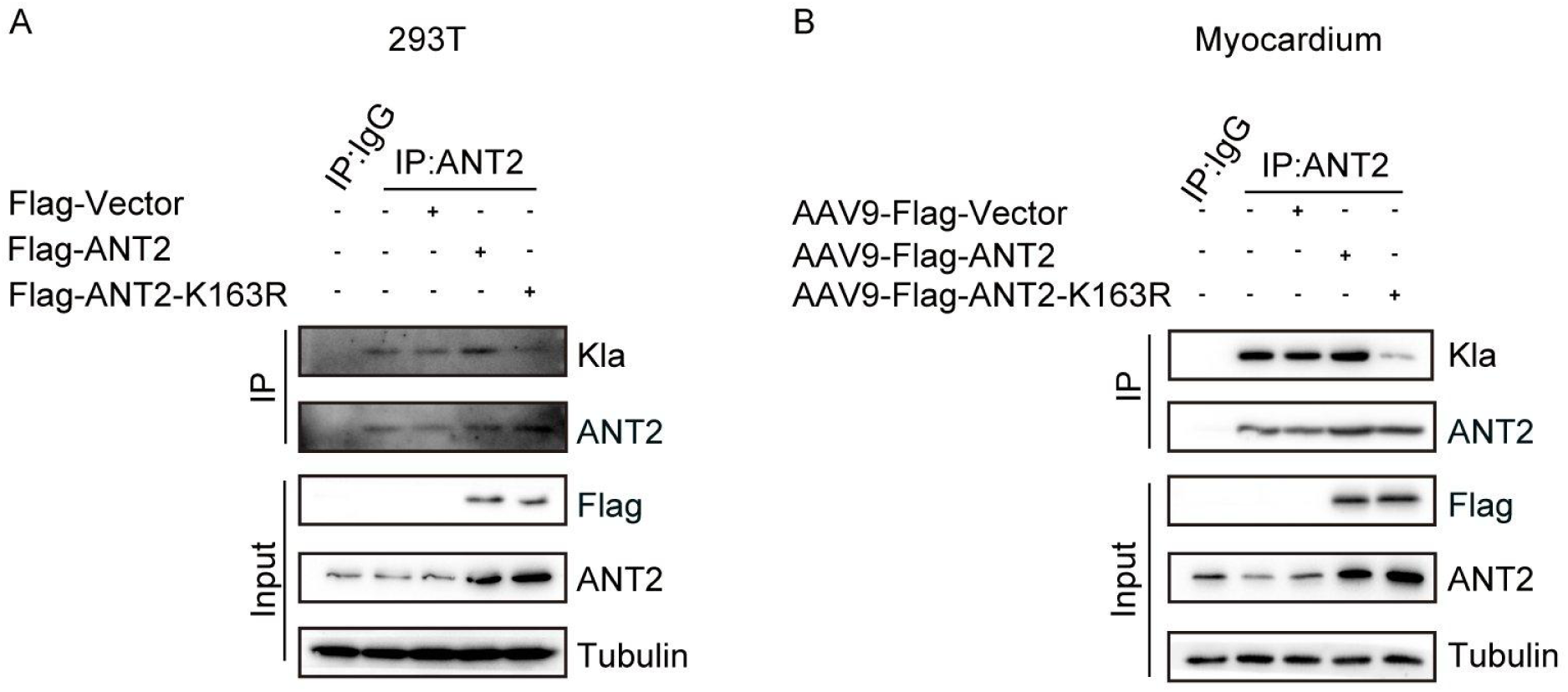
ANT2-K163R is less sensitive to lactylation modification. (**A**) Western blot analysis of lactylated ANT2 and ANT2-K163R following immunoprecipitation of ANT2 in HEK293T cells. (**B**) Western blot analysis of lactylated ANT2 and ANT2-K163R following immunoprecipitation of ANT2 in mouse myocardial tissues. Note that ANT2-K163R shows reduced lactylation compared to ANT2.

**Fig. S11.**
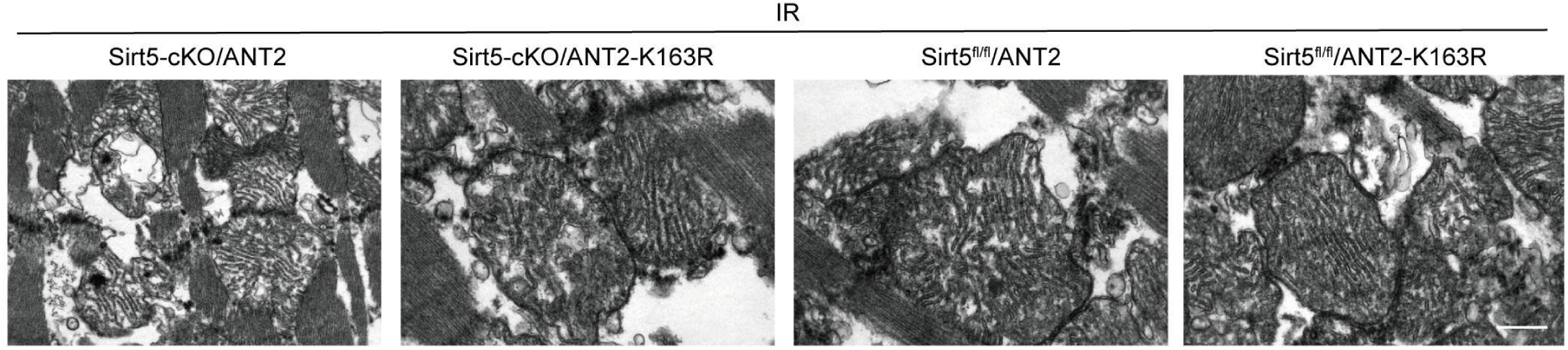
ANT2-K163R protects mitochondrial integrity in the myocardium. Overexpression of ANT2-K163R reduces the extent of mitochondrial defects induced by IR injury with or without *Sirt5* conditional knockout.

**Fig. S12.**
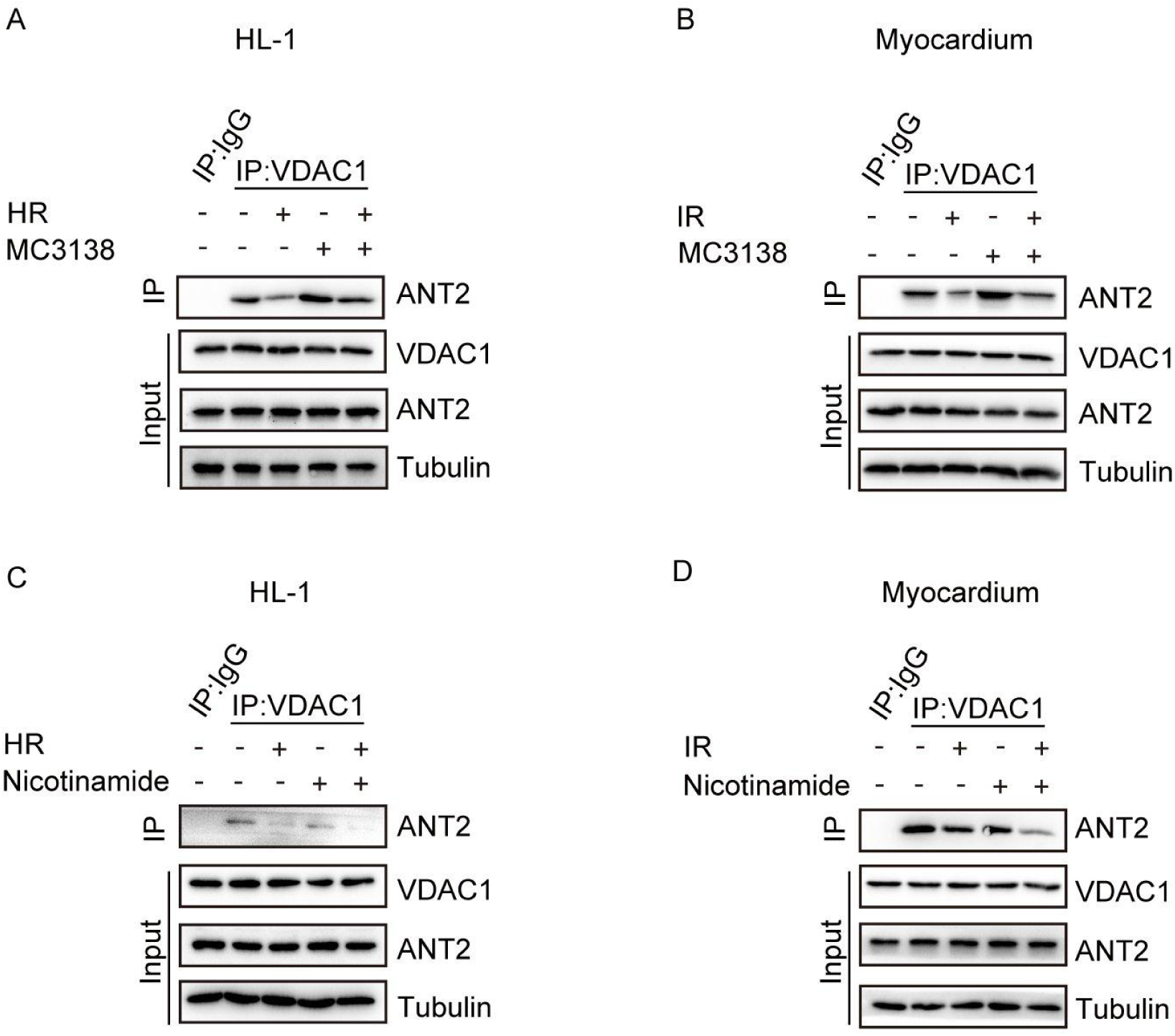
Regulation of ANT2 interaction with VDAC1 by Sirt5 activator and inhibitor. (**A** and **B**) Co-IP shows that the Sirt5 activator MC3138 promotes the interaction of ANT2 with VDAC1 in HL-1 cells and mouse myocardial tissues, which is prevented by HR or IR. (**C** and **D**) Co-IP shows that the Sirt5 inhibitor nicotinamide prevents the interaction of ANT2 with VDAC1 in HL-1 cells and mouse myocardial tissues, which is further reduced by HR or IR.

**Table S1.**
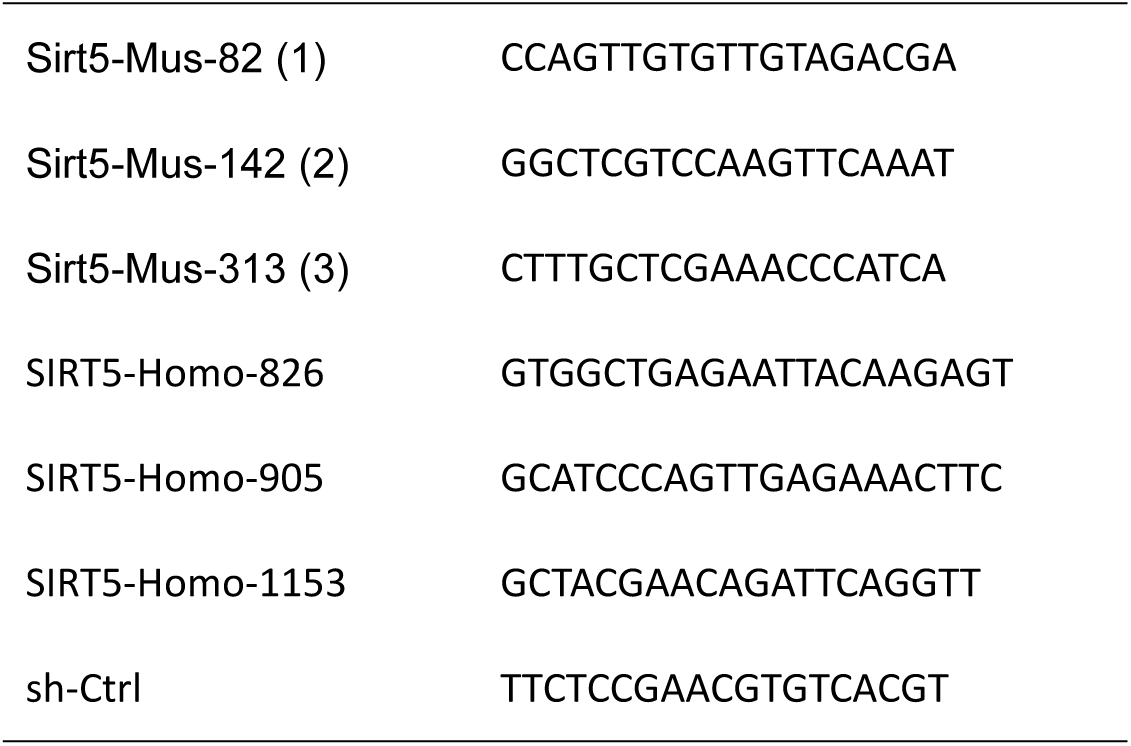
shRNA target sequences used in this study. Sirt5-Mus-82 (1) CCAGTTGTGTTGTAGACGA

**Table S2.** Antibodies used in this study.

| Antibodies | Sources | Cat. numbers | Dilutions |
| --- | --- | --- | --- |
| Ant-Sirt1 | Abcam | Ab110304 | WB (1:1000) |
| Ant-Sirt2 | Abcam | Ab134171 | WB (1:1000) |
| Ant-Sirt3 | Abcam | Ab217319 | WB (1:1000) |
| Ant-Sirt4 | Abcam | Ab231137 | WB (1:1000) |
| Ant-Sirt5 | CST | 8782S | WB (1:1000) |
|  | Thermo Fisher | PA5-20966 | IF (1:200) |
| Ant-Sirt6 | Abcam | Ab191385 | WB (1:1000) |
| Ant-Sirt7 | Abcam | Ab259968 | WB (1:1000) |
| Anti-β-tubulin | CST | 2146S | WB (1:1000) |
| Anti-L-lactyllysine rabbit mAb | PTM BIO | PTM-1401RM | WB (1:1000) |
| Anti-crotonyllysine mouse mAb | PTM BIO | PTM-502 | WB (1:1000) |
| Anti-succinyllysine mouse mAb | PTM BIO | PTM-419 | WB (1:1000) |
| Anti-β-hydroxybutyryllysine rabbit mAb | PTM BIO | PTM-1201RM | WB (1:1000) |
| Anti-acetyllysine mouse mAb | PTM BIO | PTM-101 | WB (1:1000) |
| Anti-ANT2 | Abcam | ab222843 | IP (5 μg)<br>IHC (1:100) |
|  |  | ab192410 | WB (1:500) |
| Anti-HA | Genscript Biotech | A01244 | IP (5 μg)<br>WB (1:1000) |
| Anti-Flag | Genscript Biotech | L00455B | IP (5 μg)<br>WB (1:1000) |
| Anti-VDAC1 | Abcam | ab14734 | WB (1:500) |
| HRP-conjugated affinipure goat anti-mouse IgG (H+L) | Proteintech | SA00001-1 | WB (1:10000) |
| HRP-conjugated affinipure goat anti-Rabbit IgG (H+L) | Proteintech | SA00001-2 | WB (1:10000) |

